# Acetylation-dependent remodeling of the secretory pathway shapes the senescence-associated secretome

**DOI:** 10.64898/2026.06.23.733964

**Authors:** Tornike Nasrashvili, Blenda Emini, Emilio Cirri, Norman Rahnis, Nadine Poempner, Christopher Gerner, Johannes Grillari, Regine Heller, Christoph Kaether

## Abstract

Cellular senescence is characterized by stable cell cycle arrest and the senescence-associated secretory phenotype (SASP), which drives tissue remodeling and inflammation. Underlying SASP with its increased secretion of cytokines and other secreted proteins is a massive reorganization of the secretory pathway. While transcriptional regulation of senescence has been extensively studied, the contribution of post-translational modifications (PTM) to secretory pathway regulation remains poorly understood.

Here, we combined quantitative proteomics with multi-layered PTM profiling of phosphorylation, ubiquitination and acetylation to investigate how intracellular trafficking and secretion are regulated in senescence. Using doxorubicin-induced senescence as the primary model, we identified extensive proteome remodeling, with pronounced changes in ER–Golgi-associated pathways and secretory machinery. Acetylation emerged as the most prominently regulated PTM, particularly affecting proteins involved in vesicle trafficking and ER proteostasis. Comparable proteome and PTM remodeling were also observed in replicative senescence, indicating that these changes are not restricted to a single senescence model.

Functional analyses revealed activation signatures of the acetyltransferases p300/CBP, linking global acetylation changes to enzymatic activity. Pharmacological inhibition of p300/CBP using A485 selectively modulated senescence-associated features without reversing growth arrest, consistent with a senomorphic-like effect during senescence establishment. Secretome profiling further demonstrated changes in the composition of secreted factors, consistent with modulation of the senescence-associated secretory phenotype.

Together, these findings indicate that acetylation-dependent regulation of the secretory pathway shapes the senescence-associated secretome, revealing a mechanistic link between post-translational regulation, intracellular trafficking, and extracellular signaling in senescence.

## Introduction

Cellular ageing is a multifaceted biological process driven by interacting molecular and cellular hallmarks, among which cellular senescence has emerged as a central contributor to altered tissue homeostasis and intercellular communication (Lopez-Otin et al., 2013, 2023). Senescence is commonly defined as a largely irreversible growth arrest (Hayflick & Moorhead, 1961) triggered by diverse stressors—including genotoxic injury, telomere dysfunction, and oncogenic signalling—accompanied by extensive phenotypic reprogramming at the levels of chromatin state, metabolism, organelle organisation, and proteostasis (Campisi & d’Adda di Fagagna, 2007; Kuilman et al., 2010). Although senescence can impose tumour-suppressive constraints early in life, senescent cells can accumulate and contribute to age-associated dysfunction through inflammatory and matrix-remodelling signals that act on neighbouring cells and tissues (Campisi & d’Adda di Fagagna, 2007; Coppé et al., 2010; Wang et al., 2024). This has motivated growing interest in interventions that attenuate detrimental senescence phenotypes without necessarily eliminating senescent cells. Such approaches, often termed senomorphic, aim to suppress maladaptive features such as the SASP while preserving stable growth arrest (Saliev & Singh, 2025).

A defining effector of senescence is the senescence-associated secretory phenotype (SASP), a complex hypersecretory program comprising cytokines, chemokines, growth factors, proteases, and extracellular matrix (ECM) regulators (Coppé et al., 2010; Wang et al., 2024). SASP establishment is closely linked to persistent stress signalling. For example, persistent DNA damage response signalling has been shown to trigger senescence-associated inflammatory cytokine secretion, providing a mechanistic connection between genotoxic triggers and paracrine inflammatory output (Rodier et al., 2009). The SASP is also heterogeneous across cell types and inducers, and its complexity is frequently underestimated when assessed using a limited marker panel (Basisty et al., 2020). Proteomic mapping efforts demonstrate that hundreds of proteins can be differentially secreted in each senescence context and that the secreted output cannot be reliably inferred from transcript abundance alone (Basisty et al., 2020; Coppe et al., 2008).

SASP production and export impose substantial demand on the secretory pathway, which coordinates cargo folding and quality control in the endoplasmic reticulum (ER), vesicle budding and targeting, trafficking through the Golgi, and delivery to the plasma membrane or extracellular space (Barlowe & Miller, 2013; Bonifacino & Glick, 2004; Hamazaki & Murata, 2024; Llewellyn et al., 2023; Sabath et al., 2020; Wang et al., 2024). Senescent cells must therefore adapt not only at the transcriptional level to upregulate secreted factors, but also at the level of intracellular trafficking and proteostasis to sustain chronic secretion (Coppé et al., 2010; Wang et al., 2024). However, while transcriptional regulation of senescence and SASP programs has been intensively studied, there remains a need for integrated protein-level frameworks that connect senescence-associated secretory demand to systems-level remodelling of the secretory machinery and its regulatory layers, including post-translational modifications (PTMs).

One candidate regulatory axis linking transcriptional output to secretory adaptation is centered on the lysine acetyltransferases p300/CBP, transcriptional co-activators with broad chromatin-regulatory roles that can be selectively targeted using catalytic inhibitors (Lasko et al., 2017; Yu et al., 2025). In senescence, p300 has been shown to induce de novo super-enhancers that drive senescence programs, supporting the concept that p300 activity can act upstream of stable phenotypic remodelling (Di Fede et al., 2025; Sen et al., 2019). These observations motivate the hypothesis that p300/CBP-dependent acetylation not only regulates SASP-associated gene expression at the transcriptional level but may also contribute more broadly to the cellular adaptations required to sustain secretion, including changes in intracellular organization, protein homeostasis, and trafficking capacity.

In this study, we use doxorubicin-induced stress-induced premature senescence (SIPS) in MRC-5 fibroblasts alongside complementary growth arrest and replicative senescence comparisons to define how senescence remodels the cellular proteome, and multiple PTMs, including acetylation, phosphorylation, and ubiquitination, with a particular focus on the secretory pathway. We then test functional dependence on the p300/CBP catalytic axis using the selective inhibitor A485 and quantify consequences for senescence features and proteome configuration. Finally, to directly assess SASP, we perform label-free secretome proteomics across DMSO, A485, doxorubicin, and A485+doxorubicin conditions. Together, this combined intracellular–extracellular proteomics framework connects senescence-associated acetylation programs to secretory pathway remodeling and measurable SASP secretion.

## Results

### Doxorubicin induces canonical stress-induced premature senescence (SIPS) and remodels secretory compartments

To establish a robust model of stress-induced premature senescence (SIPS), MRC-5 fibroblasts were treated with doxorubicin (DOXO) for seven days and compared to DMSO-treated controls. Senescence induction was first assessed by senescence-associated β-galactosidase (SA-β-gal) staining, which showed a strong increase in SA-β-gal–positive cells upon DOXO treatment (Figure 1A–B). In parallel, crystal violet staining revealed a pronounced reduction in cell density in DOXO-treated cells, suggesting a stable growth arrest (Figure 1C).

**Figure 1.**
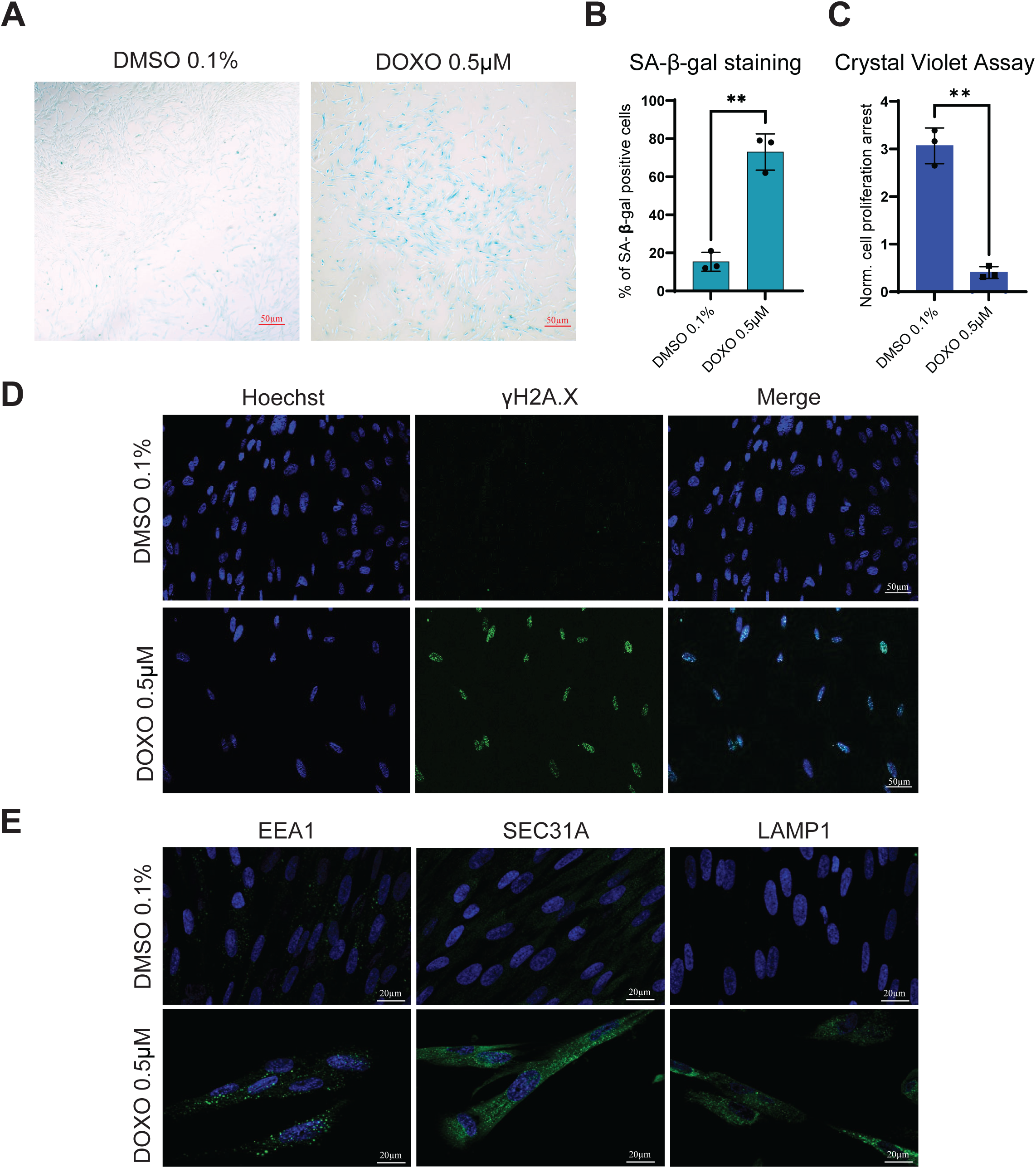
Doxorubicin Induces Senescence and Reorganizes Secretory-Pathway Compartments in MRC-5 Cells. (A) MRC-5 fibroblasts were treated with vehicle control (DMSO, 0.1%) or doxorubicin (DOXO, 0.5 µM) for 7 days, fixed, and stained for senescence-associated β-galactosidase. Representative brightfield images were acquired using an EVOS XL Core imaging system. (B) Quantification of β-galactosidase–positive cells corresponding to the images in (A). (C) Quantification of Crystal violet staining of the same culture plates, with absorbance measured at 570 nm as a proxy for cell density. (D) Immunofluorescence staining of γH2A.X to assess doxorubicin-induced DNA damage, visualized as nuclear puncta (Alexa Fluor 488). (E) Cells treated as in (A) were fixed and immunostained for EEA1 (early endosomes), LAMP1 (lysosomes), or SEC31A (ER exit sites). Secondary antibodies conjugated to Alexa Fluor 488 were used for detection. Images represent single optical sections acquired under identical imaging conditions using a Zeiss Apotome optical sectioning system. Representative images are shown. Panels A-F, n=3 independent experiments. The graph shows the mean ± standard deviation (SD). Statistical significance was assessed using unpaired t test. Significance indicators: ****p ≤ 0.0001; ***p ≤ 0.001; **p ≤ 0.01; *p ≤ 0.05; ns, not significant.

To confirm activation of a persistent DNA damage response, cells were stained for γH2AX, a marker of DNA double-strand breaks. DOXO-treated cells displayed prominent nuclear γH2AX foci compared to controls (Figure 1D), consistent with sustained genotoxic stress driving senescence (d’Adda di Fagagna et al., 2003; Rodier et al., 2009).

Given the central role of secretory activity in senescence, we next examined the organization of intracellular trafficking compartments using immunofluorescence microscopy. DOXO-treated cells exhibited increased staining intensity of EEA1, SEC31A, and LAMP1 (Figure 1E), indicating alterations in early endosomes, ER exit sites, and lysosomal compartments, respectively. ERGIC-53 staining appeared moderately increased in DOXO-treated cells, however, ERGIC-53 protein abundance was not altered in whole-cell proteomics or immunoblot analysis, suggesting that this may reflect changes in compartment organization or staining pattern rather than increased protein expression. In contrast, GIANTIN and TGN46, markers of the Golgi and trans-Golgi network, respectively, did not show obvious changes (Supplementary Figure 1). The data suggest that senescence is associated with selective remodeling of secretory pathway compartments rather than uniform reorganization of the entire trafficking network.

### SIPS Broadly Reprograms the Proteome of MRC-5 Cells

We next characterized the global proteomic changes associated with doxorubicin-induced SIPS. To this end we performed whole-cell proteomics in DMSO- and DOXO-treated MRC-5 fibroblasts cells. As an additional non-senescent growth arrest control, contact-inhibited (CI) cells were included to distinguish senescence-specific remodeling from changes associated with proliferation arrest alone (Figure 2).

**Figure 2.**
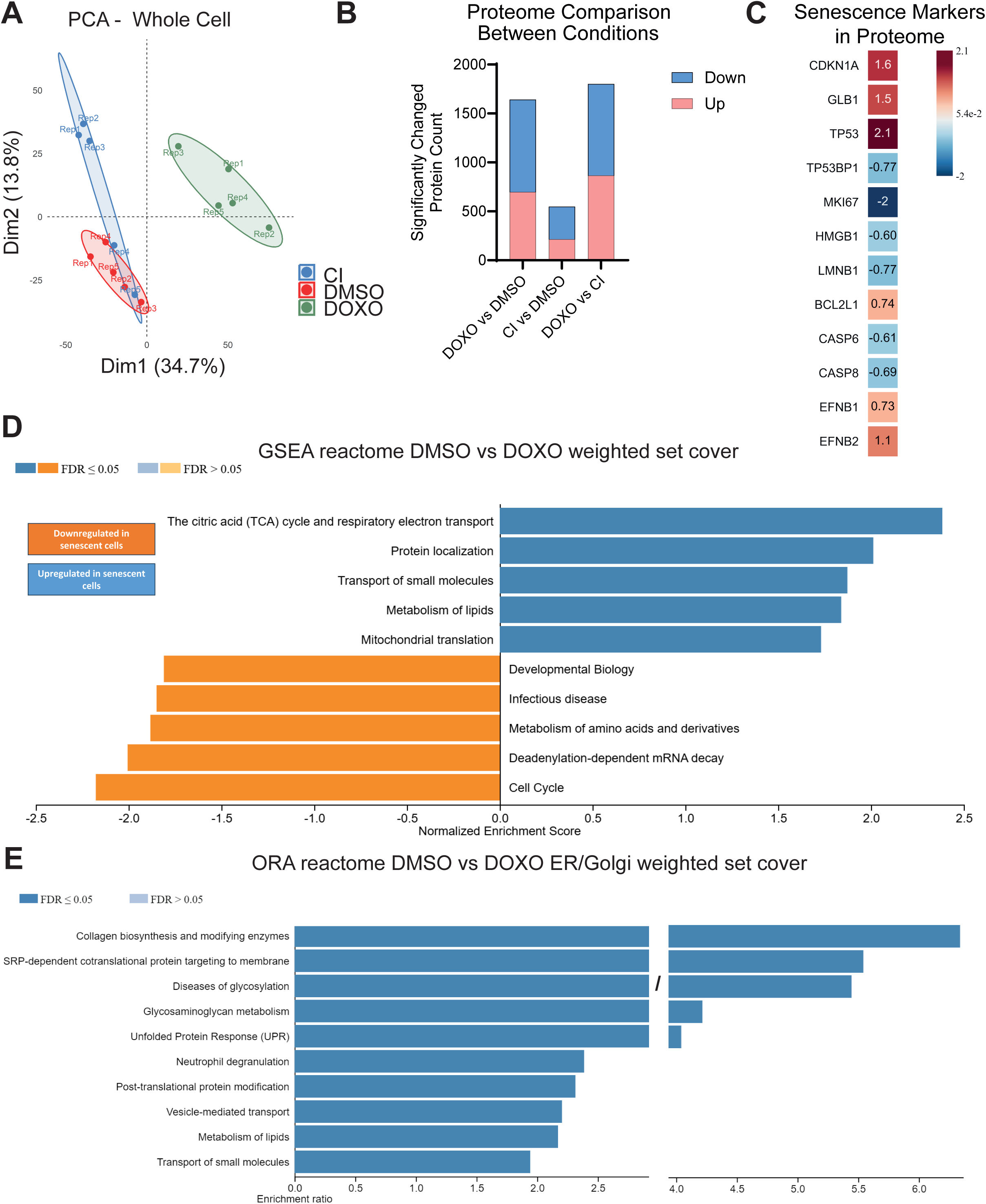
Senescence Broadly Reprograms the MRC-5 Proteome. (A) Principal component analysis (PCA) of whole-cell proteomics data (n = 5 per condition) comparing DMSO-treated controls, DOXO-induced senescent cells, and contact-inhibited (CI) cells (contact inhibition–induced cell cycle arrest). (B) Total number of significantly regulated proteins (log₂FC ≤ –0.58 or ≥ 0.58, adjusted p ≤ 0.05) between condition comparisons. (C) Heatmap depicting senescence-associated markers identified in the proteomics dataset in the comparison of DMSO versus DOXO conditions. Numbers indicate log₂FC (D) Gene set enrichment analysis (GSEA) of Reactome pathways performed with WebGestalt, showing up- and downregulated pathways in DOXO-treated cells relative to controls. (E) Over-representation analysis (ORA) of Reactome pathways performed with WebGestalt to identify functional categories altered in senescence of proteins localized in ER and Golgi.

Unsupervised principal component analysis (PCA) revealed clear separation of DOXO-treated samples from both DMSO and CI conditions (Figure 2A), indicating that SIPS is accompanied by a distinct proteomic state beyond generic cell-cycle arrest. Differential expression analysis identified a substantial number of significantly regulated proteins in DOXO-treated cells compared to controls (Figure 2B, Supplementary Table 1), consistent with widespread proteome remodeling during senescence.

Inspection of established senescence-associated markers confirmed the validity of the dataset. DOXO-treated cells displayed increased abundance of proteins linked to senescence and stress responses, including CDKN1A (p21), GLB1, and TP53, alongside decreased levels of proliferation-associated and nuclear integrity proteins such as MKI67, LMNB1, and HMGB1 (Figure 2C). These changes are consistent with stable growth arrest, lysosomal expansion, and senescence-associated nuclear remodeling (Coppe et al., 2008; Hernandez-Segura et al., 2018).

To gain functional insight into the proteomic changes, we performed pathway enrichment analyses. Gene set enrichment analysis (GSEA) revealed upregulation of metabolic pathways, including mitochondrial respiration and lipid metabolism, alongside downregulation of cell-cycle–associated processes (Figure 2D), consistent with metabolic rewiring accompanying senescence (Wiley & Campisi, 2016).

Given the focus of this study on secretory regulation, we next focused on ER- and Golgi-associated proteins. Proteins were classified based on annotated subcellular localization, and those assigned to ER or Golgi apparatus were selected for downstream analysis (Supplementary Table 2). Over-representation analysis (ORA) performed on this subset identified enrichment of pathways related to collagen biosynthesis, protein glycosylation, vesicle-mediated transport, and SRP-dependent targeting to the ER (Figure 2E). Additionally, pathways associated with the unfolded protein response (UPR) and ER stress were enriched, consistent with increased secretory demand and proteostasis burden in senescent cells (Basisty et al., 2020; Coppé et al., 2010).

To assess whether these observations extend beyond the chemically induced SIPS model, we analyzed replicative senescence in HUVECs across passages (P1, P5, P20). Late-passage cells displayed classical senescence features, including increased SA-β-gal positivity and elevated p21/p16 levels, consistent with previous reports using comparable HUVEC replicative senescence models (Stabenow et al., 2022), and were accompanied by progressive proteome remodeling and separation of early and late passage samples in PCA (Supplementary Figure 2A). However, the overall magnitude of proteomic changes was less pronounced compared to the MRC-5 SIPS model, as reflected by the lower number of significantly regulated proteins (Supplementary Figure 2C, Supplementary Table 5). Because these comparisons differ in both cell type and senescence trigger, model-specific and cell-type-specific effects cannot be fully separated. Nevertheless, the detection of shared changes despite these differences suggests that part of the response may reflect common senescence-associated regulation.

Together, these results indicate that while proteome remodeling is a conserved feature of senescence across models, the extent of these changes is context-dependent, with chemically induced SIPS exhibiting a stronger and more extensive reprogramming. Notably, enrichment of secretory and ER/Golgi-associated pathways in the MRC-5 model highlights a potential link between senescence-associated proteome remodeling and increased reliance on intracellular trafficking and proteostasis mechanisms that support SASP production.

### Multi-PTM profiling identifies acetylation as the most prominent senescence-regulated modification layer

To investigate how senescence reshapes post-translational regulation beyond protein abundance changes, we profiled acetylation, phosphorylation, and ubiquitination in the same MRC-5 samples used for whole-proteome analysis (DMSO, DOXO, and contact-inhibited (CI) controls; Figure 3A–F). This matched multi-omics framework enables discrimination between true PTM-specific regulation and changes driven by protein abundance alone.

**Figure 3.**
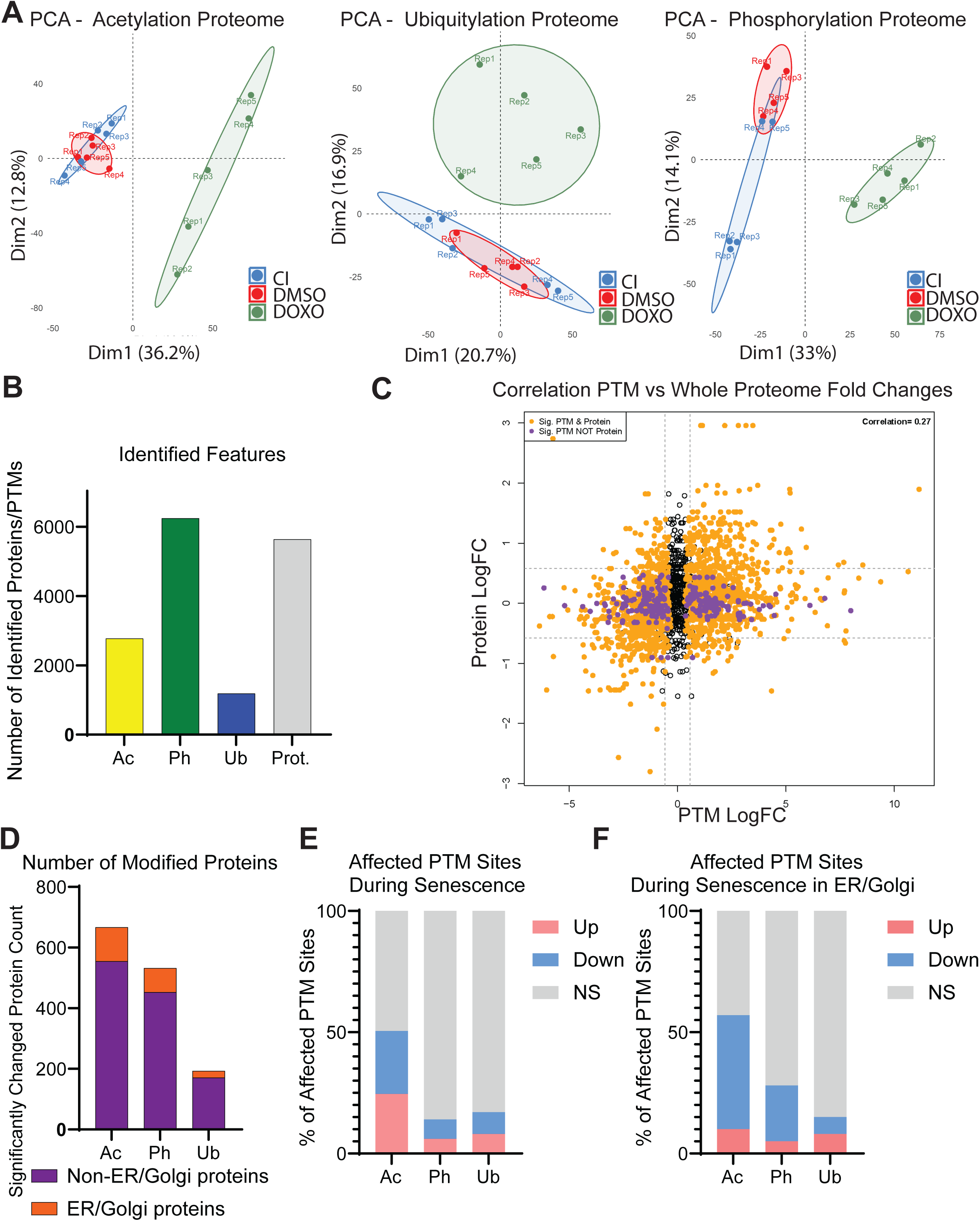
Senescence Reshapes the PTM Landscape with Strong Acetylation Changes in ER/Golgi Proteins. (A) Principal component analysis (PCA) of acetylation, phosphorylation, and ubiquitination datasets comparing DMSO, DOXO and contact-inhibited (CI, contact-inhibited) cells (n = 5 per condition). (B) Number of identified features (significant and non-significant protein signals detected by MS) in each PTM category and across the whole proteome. (C) Scatterplot describing the correlation between proteome and acetylome, in orange are highlighted proteins that are affected significantly in both layers (Ac Qvalue≤0.05; proteome adj. P.Value ≤ 0.05), in purple proteins which are significantly changing only for acetylation (Ac Qvalue≤0.05; proteome adj. P.Value > 0.05). (D) Bar chart showing distribution of significantly modified proteins into two subsets, ER and Golgi proteins in orange, Non-ER and Golgi in purple. (E) Distribution of up- and downregulated modification sites across PTMs in the whole proteome. (F) Distribution of up- and downregulated modification sites across PTMs in ER- and Golgi-associated proteins. Cutoffs for PTM enrichment analysis were log₂FC ≤ –0.58 or ≥ 0.58 with q-value ≤ 0.05.

PCA of the PTM datasets revealed clear separation of DOXO-treated samples from control conditions, whereas DMSO and CI samples showed more limited separation depending on the PTM (Figure 3A), indicating that senescence is accompanied by coordinated remodeling of multiple PTMs beyond generic cell-cycle arrest. Correlation analysis between PTM fold changes and corresponding protein abundance further demonstrated that a substantial fraction of PTM alterations occurs independently of protein-level changes (Figure 3C), supporting the presence of regulatory modification-specific programs.

Although phosphorylation yielded the highest number of identifiable features (Figure 3B), this may primarily reflect greater detection depth rather than regulatory prominence. When focusing on significantly regulated proteins, acetylation accounted for the largest number of modified proteins across the dataset (Figure 3D). Importantly, when normalized to the total number of detected sites, acetylation also exhibited the highest proportion of significantly regulated modification sites (Figure 3E, Supplementary Tables 3 and 4). This pattern was further emphasized within the ER/Golgi-associated subset, where acetylation represented the dominant regulated PTM layer and displayed a bias toward downregulated sites (Figure 3F, Supplementary Table 4).

Ubiquitination profiling also revealed condition-dependent changes across the dataset, indicating that protein turnover pathways are modulated during senescence. However, compared to acetylation and phosphorylation, ubiquitination yielded substantially fewer detectable features (Figure 3B) and significantly regulated sites (Figure 3D–E), limiting statistical power for downstream analyses. This difference was further reflected in the reduced representation of ubiquitination changes within the ER/Golgi-associated subset (Figure 3D and F). Consequently, ubiquitination was not pursued in comparative analysis with the HUVEC replicative senescence model. Nevertheless, the observed changes suggest that ubiquitin-dependent regulatory mechanisms may contribute to senescence-associated proteome remodeling and warrant dedicated investigation in future studies.

The changes in acetylation of ER/Golgi-associated proteins in senescence suggest that acetylation may contribute to the regulation of intracellular trafficking and proteostasis pathways required to sustain the SASP, which imposes a substantial burden on the secretory machinery (Basisty et al., 2020; Coppe et al., 2008; Yang et al., 2025).

To assess whether these PTM patterns extend beyond the chemically induced SIPS model, we analyzed acetylation and phosphorylation in the replicative senescence HUVEC model (P1, P5, P20). While both PTM layers showed passage-dependent remodeling, the separation between conditions and the overall magnitude of changes were less pronounced compared to the MRC-5 SIPS model (Supplementary Figure 2B,D-I, Supplementary Tables 6 and 7). Due to reduced statistical power in the HUVEC dataset, a nominal p-value threshold (p ≤ 0.05) was applied, whereas the MRC-5 DOXO dataset was analyzed using multiple-testing correction (adjusted p ≤ 0.05). Thus, while similar trends are detectable, PTM remodeling appears more robust in the chemically induced senescence context.

To further evaluate the overlap of PTM regulation across models and cell types, we compared individual acetylated sites in MRC-5 SIPS (DMSO vs DOXO) and HUVEC replicative senescence (P1 vs P20). A total of 120 proteins carried identical acetylation sites across both models (Supplementary Figure 3A). Focusing on ER/Golgi-associated proteins, 18 proteins exhibited conserved site-specific acetylation (Supplementary Figure 3B). These conserved targets include key components of the secretory pathway, encompassing proteins involved in ER–Golgi trafficking (e.g., ARF1, COPA, SEC24D, SEC61A1), protein folding and ER quality control (e.g., CANX, HSPA5, HSP90B1), as well as regulators of calcium and redox homeostasis. A full functional classification is provided in Table 1. In contrast, the overlap in identically phosphorylated sites was smaller (Supplementary Figure 3C–D).

**Table 1.**
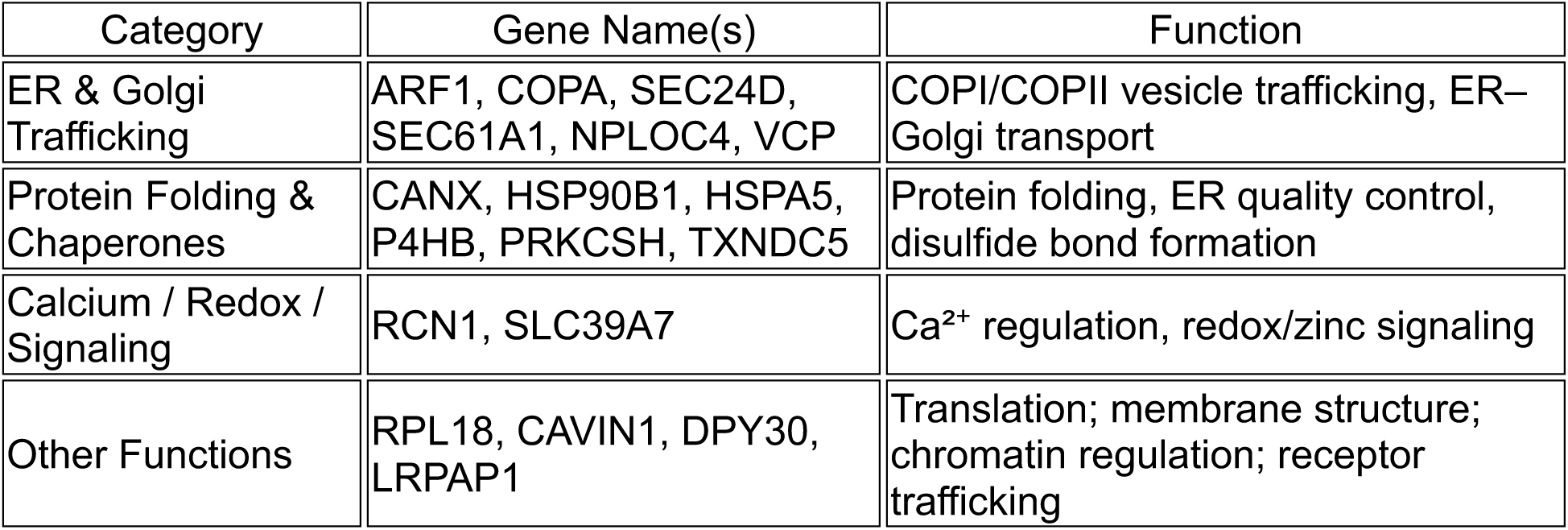
Protein (gene names) classification table of conserved acetylation sites in ER and Golgi proteins across two senescent models.

| Category | Gene Name(s) | Function |
| --- | --- | --- |
| ER & Golgi Trafficking | ARF1, COPA, SEC24D, SEC61A1, NPLOC4, VCP | COPI/COPII vesicle trafficking, ER–Golgi transport |
| Protein Folding & Chaperones | CANX, HSP90B1, HSPA5, P4HB, PRKCSH, TXNDC5 | Protein folding, ER quality control, disulfide bond formation |
| Calcium / Redox / Signaling | RCN1, SLC39A7 | Ca<sup>2+</sup> regulation, redox/zinc signaling |
| Other Functions | RPL18, CAVIN1, DPY30, LRPAP1 | Translation; membrane structure; chromatin regulation; receptor trafficking |

Functional enrichment analysis of shared PTM-regulated proteins revealed distinct biological themes. Proteins with conserved phosphorylation sites were predominantly enriched for cytoskeletal organization, whereas shared acetylation targets spanned multiple functional modules (Supplementary Figure 4), suggesting that acetylation captures a broader range of senescence-associated processes (Hwang et al., 2023; Yang et al., 2025), while phosphorylation reflects more specialized regulatory programs.

Together, these analyses identify acetylation as the most prominently remodeled PTM in senescence, particularly within ER/Golgi-associated proteins, and reveal partial but context-dependent overlap of PTM regulation across senescence models and cell types. These findings suggest that there are acetylation-dependent regulatory mechanisms, including those mediated by acetyltransferases, as potential upstream drivers of senescence-associated proteome and secretory pathway remodeling.

### Acetylation changes in secretory pathway proteins map to trafficking and ER proteostasis programs

Given that acetylation emerged as the most prominently remodeled PTM in senescence (Figure 3), we next sought to determine the biological processes associated with acetylation changes in ER/Golgi-associated proteins. To this end, we performed functional network analysis on proteins carrying significantly regulated acetylation sites within the ER/Golgi subset.

Cytoscape (ClueGO) and STRING-based enrichment analyses support a regulatory role of acetylation in proteins that cluster into distinct but functionally interconnected modules (Figure 4). As expected from the ER/Golgi-restricted protein set, vesicle-mediated transport and ER-associated proteostasis pathways, including protein folding, ER stress response, and UPR, were prominently represented.

**Figure 4.**
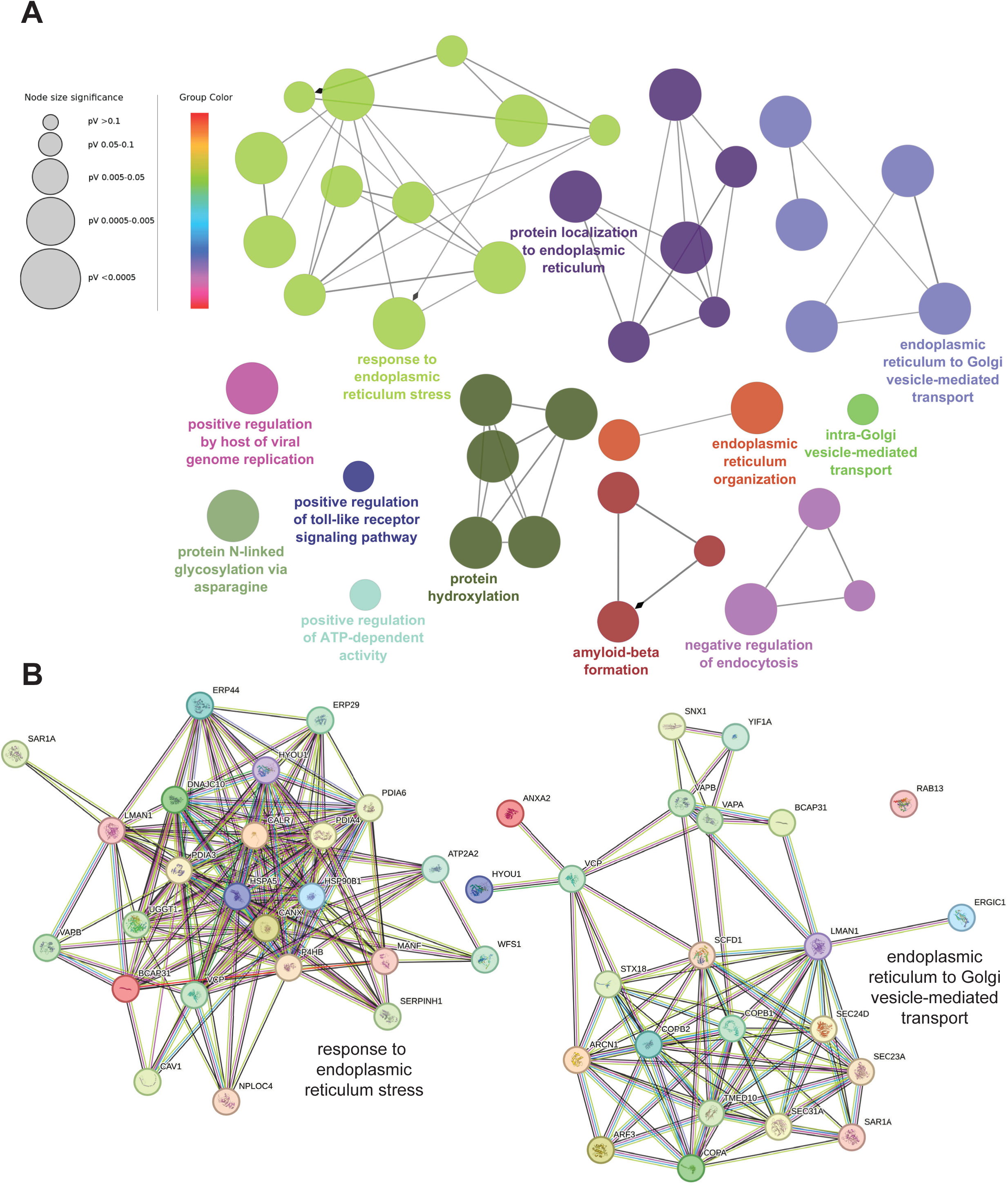
Acetylome Rewiring in Senescence Highlights ER to Golgi Trafficking Machinery and ER Stress–Related Networks. (A) Biological process network generated in Cytoscape with the ClueGO plugin, based on acetylated ER- and Golgi-associated proteins identified in DOXO–treated cells compared to DMSO-treated cells. Nodes represent functional pathways in which these proteins are involved during senescence. Node size reflects statistical significance (p-value), with larger nodes corresponding to more significant enrichment. Node color indicates functional grouping of related pathways as defined by ClueGO. (B) Visualization of two clusters (with gene names) shown in Figure 4A using STRING (https://string-db.org/ version 12.0). For the list of proteins used for pathway analysis refer to the Suppl. Table 3.

The enrichment of vesicle trafficking pathways highlights the potential involvement of acetylation in regulating intracellular transport processes required for cargo delivery and secretion. In parallel, the identification of ER stress and protein folding pathways suggests that acetylation remodeling may also impact proteostasis mechanisms that are essential for maintaining protein quality under conditions of increased biosynthetic and secretory demand.

Together, these results demonstrate that senescence-associated acetylation remodeling is enriched in ER/Golgi networks governing both vesicle trafficking and proteostasis, highlighting a potential role for acetylation in coordinating intracellular processes required for sustained secretory activity.

### Senescence is accompanied by p300/CBP activation signatures and increased histone acetylation

We next asked whether the prominent changes in acetylation are associated with changes in the activity of major lysine acetyltransferases that could plausibly underlie a global acetylation program. We focused on p300/CBP, a central acetyltransferase complex and established regulator of senescence-associated transcriptional programs, whose catalytic activity is regulated by autoacetylation within an activation loop in the HAT domain (Dancy & Cole, 2016; Thompson et al., 2004).

Acetylome quantification revealed increased acetylation of p300 at lysine residues K1542 and K1546 in DOXO-treated cells compared to DMSO controls (Supplementary Figure 5A-B). These residues lie within the activation loop (amino acids ∼1520–1560), where autoacetylation relieves autoinhibition and enhances catalytic activity (Dancy & Cole, 2016; Thompson et al., 2004). Notably, total p300 protein abundance remained unchanged, indicating that the increased acetylation of p300 reflects modification-level regulation rather than altered expression. A similar pattern was observed for CBP, which also displayed increased autoacetylation at multiple HAT domain residues without changes in protein levels, consistent with coordinated regulation of both paralogs (Supplementary Table 2).

To determine whether these activation-associated signatures of p300/CBP are reflected in downstream acetylation output, we examined canonical histone substrates of p300/CBP (Hogg et al., 2021). Acetylation of histone H3 at multiple lysine residues (K9, K14, K18, K23, and K27) was significantly increased in DOXO-treated cells (Supplementary Figure 5C), while total H3 protein levels remained unchanged. These sites are well-established targets of p300/CBP and are associated with transcriptionally active chromatin and regulation of senescence-associated gene expression programs (Dancy & Cole, 2016; Sen et al., 2019).

To assess whether similar patterns are observed across senescence models, we examined the acetylome of HUVEC replicative senescence. Autoacetylation within the corresponding p300 activation loop (K1558 and K1560) (Supplementary Table 4) was also detectable in late-passage cells, although with weaker statistical support compared to the MRC-5 SIPS model. In contrast to the fibroblast system, we did not observe a comparably strong increase in downstream histone H3 acetylation, suggesting that p300/CBP activation-associated signatures are present but less pronounced in this replicative senescence model, potentially reflecting both model- and cell-type-specific differences.

Together, these acetylome-derived readouts indicate that doxorubicin-induced senescence is accompanied by increased p300/CBP autoacetylation and elevated acetylation of canonical histone substrates. This pattern is consistent with enhanced p300/CBP catalytic output at the modification level and links the global acetylation shift observed in SIPS to an enzymatic axis centred on p300/CBP. These findings identify p300/CBP as a candidate upstream regulator of senescence-associated acetylation remodeling and provide a rationale for testing its functional contribution in subsequent experiments.

### p300/CBP inhibition reshapes senescence-associated phenotypes and establishes a distinct senescence-like state

Given that SIPS was associated with activation signatures of p300/CBP at the acetylation level (Supplementary Figure 5), we next asked whether pharmacological inhibition of this axis alters the cellular response to genotoxic stress. To this end, p300/CBP activity was inhibited using A485, a selective catalytic inhibitor targeting the acetyl-CoA binding pocket of the p300/CBP HAT domain (Lasko et al., 2017), prior to and during DOXO treatment (Figure 5).

**Figure 5.**
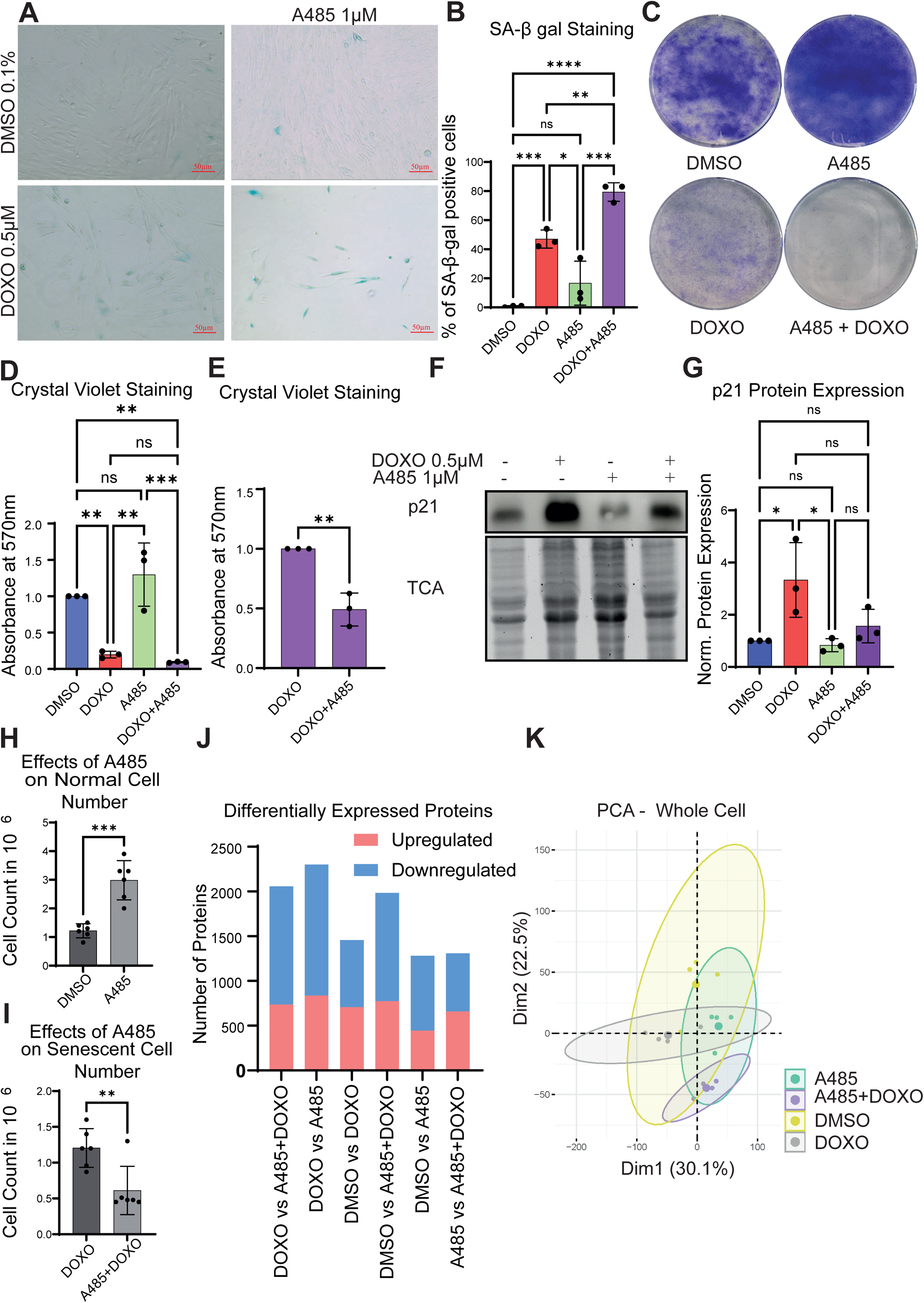
p300/CBP Inhibition with A485 Lowers p21 and Modulates Senescence Readouts in MRC-5 Cells. (A) MRC-5 fibroblasts were treated with vehicle control (DMSO, 0.1%), A485 (1 µM), doxorubicin (DOXO, 0.5 µM), or A485 + DOXO for 7 days, fixed, and stained for senescence-associated β-galactosidase. Representative brightfield images were acquired using an EVOS XL Core imaging system. (B) Quantification of SA-β-gal–positive cells corresponding to (A). (C) Representative images of crystal violet staining in four conditions: DMSO control (0.1%), A485 (1 µM), DOXO (0.5 µM), and A485 (1 µM) + DOXO (0.5 µM). (D) Crystal violet assay quantifying relative cell density (absorbance at 570 nm) across the four conditions normalized to control (DMSO 0.1%). (E) Crystal violet assay quantifying relative cell density (absorbance at 570 nm) comparison between DOXO and A485 + DOXO. Normalized to DOXO. (F) Lysates from MRC-5 fibroblasts treated with DMSO, A485, DOXO, or A485 + DOXO were subjected to SDS–PAGE and Western blot. Blots were probed with an antibody against p21; stain-free total protein detection with 10% TCA served as the loading control. (G) Quantification of p21 protein expression normalized to DMSO control. (H) Quantification of cell number in normally cycling MRC-5 cells treated with DMSO or A485 (1 µM) for 7–8 days. Equal numbers of cells were seeded, and the total cell count was measured at the endpoint. (I) Quantification of total cell number in doxorubicin-induced senescent cells (DOXO, 0.5 µM) treated with or without A485 (1 µM) for 7–8 days. (J) Number of differentially expressed proteins identified by label-free proteomics for the indicated comparisons. Proteins were classified as upregulated (log2FC ≥ 0.58) or downregulated (log2FC ≤ −0.58) using MSStats with a significance threshold of p ≤ 0.05. (K) Principal component analysis (PCA) of whole-cell proteomics showing distribution and separation of the four conditions (DMSO, A485, DOXO, A485+DOXO). Panels B–G, n=3, panels H-I, n=6 independent experiments. The graphs show the mean + standard deviation (SD). Statistical significance was assessed using one-way ANOVA (Panels B, D, and G) and an unpaired t test (Panel E, H, and I). Significance indicators: ****p ≤ 0.0001; ***p ≤ 0.001; **p ≤ 0.01; *p ≤ 0.05; ns, not significant. A485 is a selective inhibitor of the acetyltransferase p300/CBP.

SA-β-gal staining confirmed robust induction of senescence in DOXO-treated cells compared to DMSO controls (Figure 5A–C). Notably, co-treatment with A485 did not reduce SA-β-gal positivity, indicating that p300/CBP inhibition does not prevent the establishment of a senescence-associated phenotype.

Crystal violet staining revealed a reduction in cell density upon DOXO treatment, as expected, with a further decrease observed in the A485+DOXO condition (Figure 5D). To more directly assess cell number changes, we performed a paired seeding assay in which DMSO and A485-treated cells, as well as DOXO and A485+DOXO-treated cells, were seeded at equal densities (Figure 5H–I). Under these conditions, A485 treatment increased cell number in proliferating cells after 7 days, whereas in the context of DOXO treatment, A485 resulted in a further reduction in final cell number compared to DOXO alone. These findings indicate that p300/CBP inhibition exerts opposing effects depending on cellular context, enhancing proliferation under basal conditions while reducing it following genotoxic stress. One possible explanation is that A485-treated hyper-proliferating cells (Figure 5I) are more vulnerable to DOXO, whose DNA-damaging effects preferentially affect cycling cells (Oyeyode et al., 2025).

We next examined canonical senescence markers at the protein level. DOXO treatment induced CDKN1A (p21) expression, consistent with activation of the DNA damage response–associated senescence program (Gorgoulis et al., 2019; Kuilman et al., 2010) (Figure 5F–G). In contrast, co-treatment with A485 attenuated p21 levels relative to DOXO alone. Despite this reduction, SA-β-gal staining remained elevated and proteome-wide analysis did not indicate restoration of a proliferative state, suggesting that p300/CBP inhibition modulates specific components of the senescence program without reversing the overall senescence-associated phenotype.

To further characterize the global impact of p300/CBP inhibition, we performed whole-cell proteomics across all four conditions. Differential expression analysis revealed substantial remodeling of the proteome in response to A485, both in the presence and absence of DOXO (Figure 5J). PCA demonstrated clear separation of all four conditions, with the A485+DOXO samples forming a distinct cluster rather than overlapping with either DOXO or control conditions (Figure 5K). This indicates that p300/CBP inhibition does not simply revert the senescence-associated proteomic state but instead results in a proteomic state distinct from both DOXO-treated senescent cells and untreated controls.

To explore whether altered cell number was associated with changes in cell-cycle or apoptotic pathways, we examined selected regulators in the proteomics dataset. A485 altered CDK4 and CDK6 abundance in a context-dependent manner and increased CASP3 and BAX levels in selected conditions. However, PARP cleavage was not detected at the endpoint, suggesting that sustained executioner apoptosis was not ongoing in the remaining cells (Supplementary Figure 6A–E). Transient toxicity during the treatment period cannot be excluded, but these data do not support sustained apoptosis at the final time point.

Together, these results demonstrate that p300/CBP inhibition reshapes the cellular response to DOXO-induced stress without preventing the establishment of senescence-associated features. Instead, A485 modifies the senescence-associated state induced by DOXO, resulting in altered regulation of canonical markers, context-dependent effects on cell accumulation, and global proteome remodeling. These findings support a model in which p300/CBP activity contributes to defining the qualitative features and heterogeneity of the senescent phenotype rather than acting as a binary determinant of senescence induction.

### p300/CBP inhibition triggers compensatory remodeling of acetylation regulators and global lysine acetylation output under DOXO stress

Given the strong, context-dependent effects of A485 on growth and senescence-associated phenotypes, we next asked how p300/CBP inhibition reshapes the broader acetylation landscape under DOXO stress. Beyond p300/CBP itself, acetylation output is determined by the balance of multiple lysine acetyltransferases (KATs) and deacetylases (HDACs/sirtuins). We therefore examined whether A485 treatment is accompanied by compensatory shifts in these regulators on a proteome level.

Proteomics-based quantification indicated that A485 alters the abundance of multiple acetylation regulators, consistent with a broader adaptive response of the acetylation machinery rather than an isolated block of p300/CBP activity (Supplementary Figure 7A). In DOXO and A485-treated cells, these changes suggested partial restoration of acetylation capacity through compensatory rewiring of KAT/HDAC networks, in line with the distinct endpoint state observed in the global proteome.

To get a simple readout of overall acetylation output, we blotted for pan–acetyl-lysine. The most obvious change was a strong band around ∼15 kDa (Supplementary Figure 7B-C). While proteins in this molecular weight range may include histone species, band size alone does not allow definitive identification. Rather than reflecting global histone acetylation changes, this pattern likely represents alterations in a subset of low–molecular weight acetylated proteins, indicating that acetylation remodeling in response to DOXO and A485 is not uniform across all substrates but may differentially affect specific protein classes.

Together, these findings indicate that A485 does not simply cause a uniform decrease in acetylation output but instead reshapes the acetylation landscape under DOXO stress. This supports the idea that p300/CBP inhibition induces broader compensatory remodeling of acetylation-regulating pathways under DOXO stress.

### Pathway-level analyses suggest attenuation of ER stress programs under p300/CBP inhibition in SIPS

To pinpoint the ER/Golgi programs that are induced in senescence but also actively counteracted by p300/CBP inhibition, we used a two-step, direction-aware filtering strategy. We first defined senescence-regulated proteins by comparing DOXO vs DMSO and then intersected this set with proteins altered by A485 within the senescent background (A485+DOXO vs DOXO), applying the requirement that changes must be antagonistic across the two comparisons - i.e., proteins up in DOXO vs DMSO must be down in A485+DOXO vs DOXO, and vice versa. The resulting antagonistically regulated protein sets were then analyzed in STRING to identify enriched biological processes (Supplementary Figure 8A–B).

STRING enrichment of proteins that were senescence-induced (up in DOXO vs DMSO) but reversed by A485 (down in A485+DOXO vs DOXO) highlighted a coherent ER/Golgi-centered module. Enriched terms included response to ER stress and protein folding, alongside multiple processes related to collagen/ECM handling such as extracellular matrix organization, collagen fibril organization, and hydroxylation-related pathways (e.g., protein hydroxylation and related peptidyl-residue modification terms) (Supplementary Figure 8A). This pattern suggests that p300/CBP inhibition counter-regulates a senescence-associated ER/Golgi stress and ECM-processing network, consistent with previous studies linking senescence to SASP-associated secretome remodeling, proteostasis changes, and p300-dependent senescence programs (Basisty et al., 2020; Di Fede et al., 2025; L’Hote et al., 2022).

On the contrary, proteins that were reduced in senescence (down in DOXO vs DMSO) but restored by A485 (up in A485+DOXO vs DOXO) were enriched for broader proteostasis-linked functions including protein folding, protein metabolic process, and processes connected to cellular localization and regulation of catalytic activity/molecular function (Supplementary Figure 8B). Together, these opposing enrichment patterns are consistent with A485 not simply “turning off” senescence, but rewiring the senescent ER/Golgi proteostasis landscape by reversing specific modules induced in the DOXO state while restoring others that are suppressed.

Using an antagonistic filter strategy across DOXO-induced senescence and A485 modulation within senescence, we identified a specific ER/Golgi network dominated by ER stress, protein folding, and ECM-processing pathways. This network is induced during SIPS and selectively counter-regulated by p300/CBP inhibition.

### A485 demonstrates senomorphic-like effects by suppressing SASP production and secretion while remodeling secretory machinery

To determine whether p300/CBP inhibition affects the SASP, we performed label-free secretome analysis across all four conditions (DMSO, A485, DOXO, and A485+DOXO) matched with total cellular proteomics (Figure 6, Supplementary Figure 9). To enable direct comparison between intracellular protein abundance and secretion, whole-cell proteomics was performed under the same conditions as the secretome analysis (DMSO, DOXO, A485, and A485+DOXO). PCA of the secretome revealed clear separation of all four conditions, with A485+DOXO forming a distinct cluster rather than reverting toward either control or DOXO states (Figure 6B). Doxorubicin-induced senescence (DOXO) was characterized by a substantial increase in secreted proteins, whereas co-treatment with A485 markedly reduced this output (Figure 6A). Notably, the A485+DOXO condition showed a pronounced shift toward downregulated secreted proteins compared to DOXO alone, indicating a broad attenuation of SASP secretion. A complete list of differentially secreted proteins and matched intracellular proteome measurements is provided in Supplementary Tables 11 and 12.

**Figure 6.**
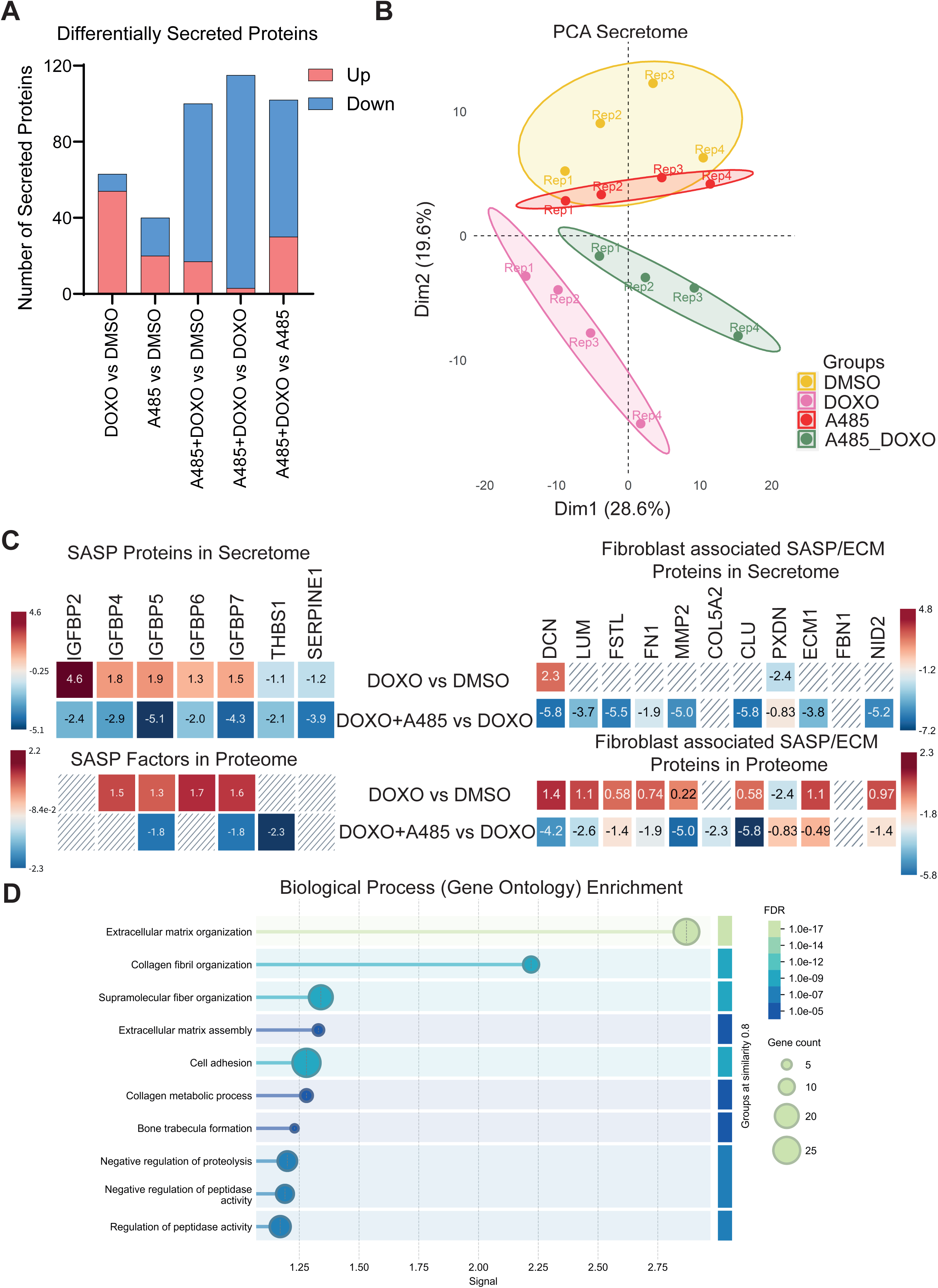
p300/CBP Inhibitor A485 Displays Senomorphic Properties and Modulates SASP-associated ER and Golgi Proteins. (A) Number of differentially secreted proteins identified by label-free secretome proteomics for the indicated comparisons. Proteins were classified as upregulated (log2FC ≥ 0.58) or downregulated (log2FC ≤ −0.58) with a significance threshold of Adj. p ≤ 0.05. Red indicates increased abundance, and blue indicates decreased abundance relative to the respective comparison. (B) Principal component analysis (PCA) of secretome showing distribution and separation of the four conditions (DMSO, A485, DOXO, A485+DOXO). (C) Heatmap showing differential expressions and secretions of canonical SASP and fibroblast specific/ECM proteins across all treatment conditions (DMSO, A485, DOXO, A485+DOXO), generated from proteomics data. Visualization created with BioRender. Numbers indicate log2fold. Hatched boxes indicate proteins that were not detected in the respective condition and therefore could not be quantified. The higher frequency of non-detected ECM proteins in the DOXO vs DMSO comparison reflects low baseline secretion levels in control cells rather than technical limitations of the dataset. Upper heatmaps show secretome data, whereas lower heatmaps show matched whole-cell proteome data for the indicated comparisons. (D) Gene ontology analysis of biological processes for secreted proteins downregulated in the A485+DOXO condition compared to DOXO alone. Quantitative values underlying this figure are provided in Supplementary Tables 11 (whole-cell proteome) and 12 (secretome). Statistical significance was assessed using one-way ANOVA. Significance indicators: ****p ≤ 0.0001; ***p ≤ 0.001; **p ≤ 0.01; *p ≤ 0.05; ns, not significant. N=4

**Figure 7.**
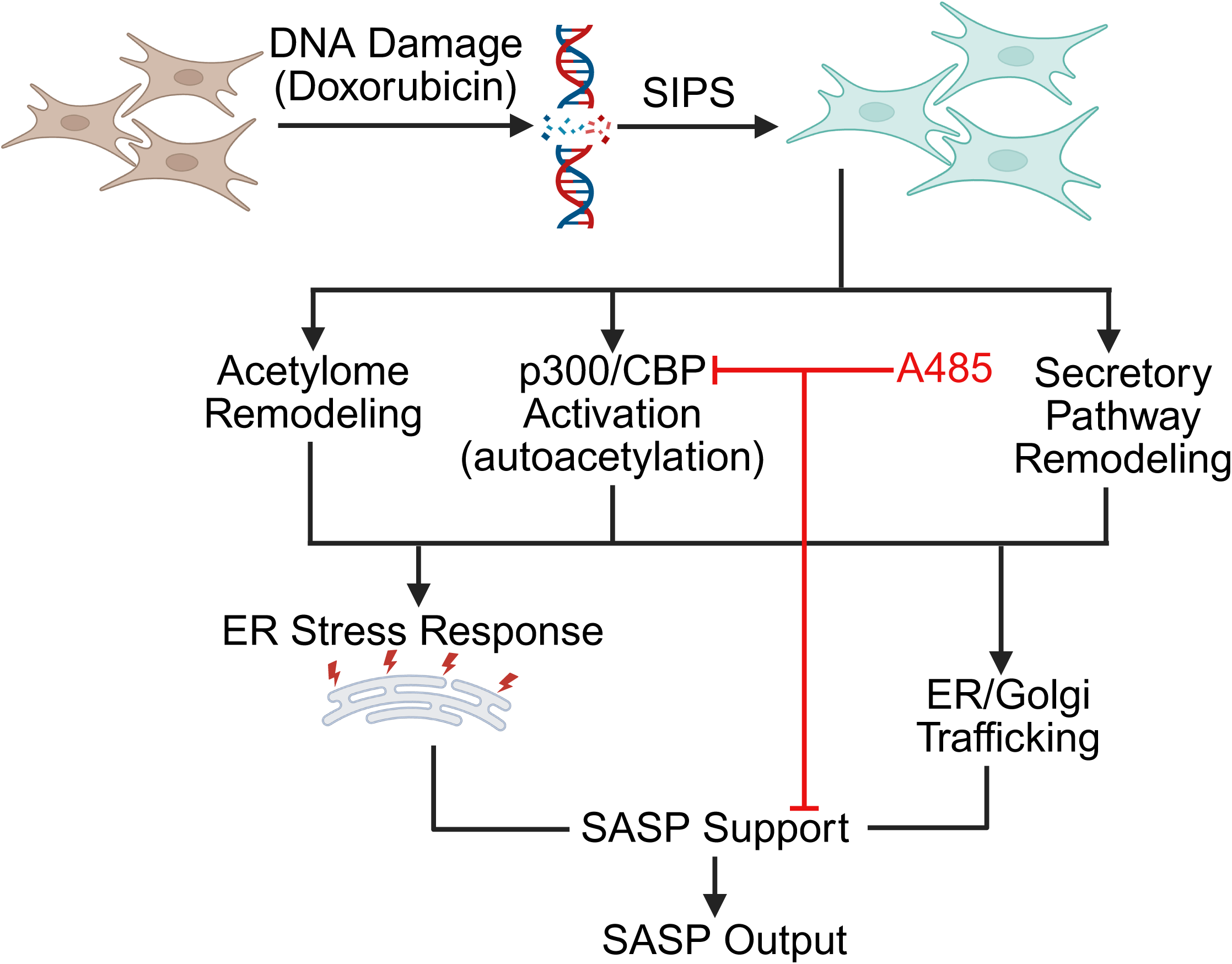
Graphical Abstract. Acetylation-dependent remodeling of the secretory pathway shapes the senescence-associated secretome. Schematic model of the proposed mechanism. Doxorubicin-induced senescence is associated with remodeling of ER/Golgi proteostasis and trafficking pathways, accompanied by prominent acetylation changes in secretory pathway proteins. p300/CBP inhibition with A485 modulates this response by attenuating SASP-associated intracellular protein abundance and secretome output without preventing senescence establishment.

This indicates that p300/CBP inhibition does not simply suppress secretion globally but instead establishes a distinct secretory phenotype.

To examine the impact on specific SASP components, we analyzed canonical SASP factors alongside fibroblast-associated extracellular matrix (ECM) proteins in both the secretome and cellular proteome. DOXO treatment led to consistent upregulation of canonical SASP factors at both the secretome and intracellular proteome levels, whereas fibroblast-associated ECM proteins were predominantly increased at the proteome level (Figure 6C). In contrast, these factors were broadly reduced in the A485+DOXO condition. Importantly, this pattern was mirrored at the intracellular proteome level, indicating that p300/CBP inhibition affects not only secretion but also the abundance of SASP-associated proteins (Figure 6C, Supplementary Figure 9B-C). These observations are consistent with the concept that the SASP represents a complex and dynamically regulated secretory program controlled at multiple levels rather than a purely secretion-driven process (Basisty et al., 2020; Wang et al., 2024). Classical pro-inflammatory cytokines were not prominently detected in the secretome dataset, which may reflect the fibroblast model, the specific DOXO-induced senescence state, or limited MS sensitivity for low-abundance cytokines. Nevertheless, multiple SASP-associated and ECM-remodeling proteins were robustly detected and modulated by A485.

Gene ontology analysis of secreted proteins downregulated in the A485+DOXO condition further supported this model, revealing enrichment of biological processes related to ECM organization, collagen fibril organization, and cell adhesion (Figure 6D). Importantly, similar biological processes were also downregulated in the A485+DOXO condition compared to A485 alone (Supplementary Figure 9D), indicating that these effects are not broadly induced by p300/CBP inhibition in proliferating cells but are instead specific to the senescent context. This specificity suggests that A485 selectively targets senescence-associated programs, including ECM remodeling processes that support SASP establishment and maintenance (Wang et al., 2024). Thus, A485 appears to shift the composition of the senescent secretome rather than abolishing secretion globally, with strongest effects on matrix-remodeling and tissue-remodeling arms of the SASP.

Together, these findings indicate that p300/CBP inhibition during senescence induction attenuates SASP-associated output without preventing the establishment of senescence-associated features. Mechanistically, this effect operates at multiple levels: (i) reduction of intracellular SASP-associated protein abundance, (ii) suppression of secretory pathway components required for cargo trafficking, and (iii) consequent decrease in extracellular SASP factors. This multi-layered regulation suggests that p300/CBP contributes to linking acetylation-dependent programs with the secretory machinery that supports the senescent phenotype (Sen et al., 2019).

### An interactive senescence PTM atlas

To make the large-scale datasets from this study more accessible, we developed an interactive senescence PTM atlas, which allows users to explore protein abundance and PTM changes across the senescence models and conditions analyzed here: https://genome.leibniz-fli.de/senescence-ptm-atlas/.

## Discussion

Cellular senescence is a multifaceted stress response characterized by stable cell cycle arrest accompanied by extensive molecular and functional reprogramming (Gorgoulis et al., 2019). In addition to its well-established tumor-suppressive role, senescence is defined by the acquisition of a SASP, which enables senescent cells to influence their microenvironment through the release of signaling molecules (Coppé et al., 2010). While transcriptional control of the SASP has been extensively studied, the contribution of post-translational regulation to secretory pathway function remains less well understood.

In this study, the use of both genotoxic stress–induced and replicative senescence models revealed a conserved program of proteome remodeling that extends beyond canonical cell cycle regulators to include pathways governing intracellular trafficking and proteostasis. These findings are consistent with previous work demonstrating that senescent cells undergo large-scale reorganization of protein networks to sustain survival under chronic stress conditions (Basisty et al., 2020; Sabath et al., 2020). Notably, the enrichment of ER- and Golgi-associated proteins highlights the central role of the secretory pathway in shaping the senescent phenotype.

A key finding of this work is the pronounced reprogramming of the post-translational modification landscape, with acetylation emerging as the most prominently regulated layer. While phosphorylation-driven signaling is critical for senescence initiation, the observed enrichment of acetylation changes in ER–Golgi-associated proteins supports a broader regulatory role in maintaining the senescent state. This is consistent with previous studies showing that acetylation and epigenetic regulation contribute to aging biology, senescence-associated transcriptional remodeling, and SASP control (Basisty et al., 2020; Benayoun et al., 2015; Sen et al., 2019; Wang et al., 2024; Yang et al., 2025). These results indicate that acetylation-dependent mechanisms extend beyond chromatin regulation and may influence intracellular trafficking machinery, consistent with emerging evidence linking lysine acetylation to cytoskeletal organization and vesicle transport (Choudhary et al., 2014; Janke & Magiera, 2020; L’Hote et al., 2022; Sen et al., 2019; Xu et al., 2025). Ubiquitination was also affected; however, it showed fewer significant and less consistent changes compared to acetylation and phosphorylation and was therefore not analyzed further. This is in line with previous work showing that ubiquitination in senescence primarily regulates protein stability and turnover, whereas acetylation more directly shapes transcriptional and functional programs, including the SASP, with both modifications acting in a coordinated but distinct manner (Yu et al., 2025).

Together, these findings suggest that acetylation changes in senescence are linked to ER/Golgi pathways involved in vesicle trafficking, protein folding, and ER stress. This supports the idea that acetylation may help coordinate intracellular protein processing and trafficking during SASP production.

Consistent with these molecular changes, senescence is associated with broad alterations of ER–Golgi networks and trafficking-related pathways. This reorganization likely reflects the increased secretory demand imposed by the SASP, which requires coordinated adaptation of protein folding, processing, and transport systems (Basisty et al., 2020; Coppé et al., 2010). The concomitant activation of ER stress–related pathways indicates that senescent cells adapt to the elevated proteostatic load. Our findings suggest that the secretory machinery is extensively reprogrammed, with functional consequences for the regulation and composition of protein secretion.

The functional relevance of acetylation-dependent regulation is further supported by the activation signatures of p300/CBP. Increased autoacetylation of these acetyltransferases, together with elevated acetylation of canonical substrates, is consistent with enhanced enzymatic activity and links global acetylation remodeling to a defined regulatory axis (Dancy & Cole, 2016; Sen et al., 2019). Notably, these activation-associated signatures occurred within a broader bidirectional remodeling of the acetylome, as senescence was characterized by both increased and decreased acetylation events overall, with ER/Golgi-associated proteins showing a relative bias toward reduced acetylation. This indicates that p300/CBP activation is likely one component of a wider regulatory network rather than the sole driver of acetylome remodeling. Although the upstream trigger of p300/CBP autoacetylation was not directly tested here, senescence-associated stress signaling and metabolic rewiring may contribute to increased acetylation capacity and p300/CBP recruitment to senescence-associated transcriptional programs. Recent preprint evidence further suggests that mitochondrial citrate and acetyl-CoA metabolism can fuel histone acetylation at SASP-associated loci and modulate SASP output (Martini et al., 2024). While p300/CBP may contribute upstream through transcriptional control of senescence programs, the compartment-specific decrease in acetylation observed in secretory pathway proteins suggests that additional acetylation-regulating enzymes may shape these changes locally. Indeed, dedicated acetyltransferase systems have been described in the ER/ERGIC, and deacetylase activity has been reported in the Golgi lumen, where acetylation can regulate trafficking and quality control of secretory cargo proteins (Farrugia & Puglielli, 2018; Pehar & Puglielli, 2013; Peng & Puglielli, 2016). Pharmacological inhibition of p300/CBP modulates key features of the senescent phenotype without abolishing growth arrest, suggesting a senomorphic role in shaping intracellular organization and stress responses (Lasko et al., 2017; Saliev & Singh, 2025; Yang et al., 2025). Our data extend this concept by showing that p300/CBP inhibition counter-regulates defined ER/Golgi-associated modules, including protein folding, ER stress, ECM-processing pathways, and SASP-associated secretome components. These observations position p300/CBP as a candidate upstream regulator of senescence-associated programs that may indirectly influence acetylation-dependent remodeling of the secretory pathway. While not necessarily responsible for all local acetylation changes within secretory compartments, p300/CBP was a logical candidate to test because of its established role in senescence and its activation signatures in our dataset.

A central mechanistic axis linking these observations is ER stress signaling. Persistent activation of UPR pathways has been implicated in both the establishment and maintenance of senescence (Hamazaki & Murata, 2024; Hetz & Saxena, 2017; Rodier & Campisi, 2011; Sabath et al., 2020). The coupling between acetylation changes and ER stress–associated networks observed here suggests that post-translational regulation contributes to coordinating adaptive responses to increased secretory and proteostatic demand. In senescent cells, moderate UPR activation is likely beneficial by supporting ER folding capacity and sustained SASP production. Prolonged activation of these pathways may simultaneously reinforce the senescent state through chronic stress signaling.

Importantly, integration of secretome analysis provides a functional link between intracellular remodeling and extracellular output. The SASP is a major driver of the non-cell-autonomous effects of senescent cells, influencing inflammation, tissue remodeling, and paracrine signaling (Coppé et al., 2010). The observed changes in secreted protein composition demonstrate that remodeling of the secretory pathway is directly translated into altered intercellular communication. The findings point to acetylation-dependent regulation of intracellular trafficking machinery as a contributor to the composition of the secretome, rather than simply controlling secretion rate.

The phenotypic effects of A485 are consistent with senomorphic-like modulations during senescence establishment, in which deleterious features of senescent cells are attenuated without reversing stable growth arrest (Lasko et al., 2017). Because A485 was applied prior to and during DOXO treatment, these data support a role for p300/CBP in shaping SASP acquisition, but do not establish whether p300/CBP inhibition can remodel an already established senescent state. In our system, p300/CBP inhibition reduced secretory and proteostasis-associated programs while preserving the arrested state, suggesting that selective modulation of the SASP may be achievable without requiring senescent cell elimination. Such an approach could be advantageous in settings where complete removal of senescent cells is undesirable, for example during wound repair or tissue regeneration, where transient senescence can exert beneficial functions (Gorgoulis et al., 2019; Hernandez-Segura et al., 2018). At the same time, p300/CBP are pleiotropic chromatin regulators involved in normal transcriptional control, differentiation, and stress responses, indicating that systemic inhibition may carry risks including impaired tissue homeostasis or reduced regenerative capacity (Martire et al., 2020). Although this remains to be tested in senescence-specific in vivo models, A485 has been reported to reduce inflammatory and tissue-remodeling programs in vivo, including LPS/D-galactosamine-induced liver injury and psoriasis-like skin inflammation (Kim et al., 2023; Peng et al., 2019). These considerations suggest that context-dependent or transient targeting strategies may be required if p300/CBP-directed senomorphic interventions are pursued therapeutically.

Several limitations of the present study should be considered. First, our multi-omics analyses identified strong associations between acetylation remodeling and secretory pathway function. However, direct validation of individual acetylation events on candidate trafficking or proteostasis proteins proved experimentally challenging and has to be addressed in future experiments. The large number of regulated targets made prioritization difficult. In addition, protein localization, stoichiometry, and function within the secretory pathway are highly cell-state dependent, complicating targeted mutation strategies. Second, the mechanistic analyses were performed primarily in a single doxorubicin-induced fibroblast model of stress-induced premature senescence, and broader validation across additional cell types and senescence inducers will be important to establish generality. In addition, comparison with the HUVEC replicative senescence dataset changes both cell type and senescence trigger, limiting the ability to distinguish model-specific from cell-type-specific effects. Third, we used only one p300/CBP inhibitor (A485). Although A485 is reported to be a selective catalytic inhibitor (Lasko et al., 2017), future studies using complementary genetic approaches or independent inhibitors would strengthen the conclusion that the observed phenotype reflects p300/CBP pathway inhibition across contexts. Furthermore, A485 was applied during senescence induction; therefore, future experiments in which p300/CBP inhibition is applied after stable SIPS establishment will be required to determine whether this axis can remodel an already established senescent state. Finally, although the observed A485 phenotype is consistent with senomorphic activity, the physiological relevance, durability, and potential trade-offs of such modulation remain to be determined in more complex tissue and in vivo settings.

Collectively, these findings indicate that senescence-associated remodeling of the secretory pathway is driven by coordinated changes in protein abundance, acetylation-dependent regulation, and sustained ER stress responses. Within this framework, acetylation emerges as a prominent regulatory mechanism that links intracellular trafficking and proteostasis to extracellular signaling outputs (L’Hote et al., 2022; Xu et al., 2025). Importantly, our data indicate that acetylation-dependent remodeling of the secretory pathway is directly coupled to and contributes to shaping the composition of the senescence-associated secretome. This identifies post-translational regulation of intracellular trafficking as a key mechanism by which senescent cells actively control their communication with the microenvironment.

### Experimental Procedures

#### Cell culture and treatments

Human fetal lung fibroblasts (MRC-5; ATCC CCL-171) were cultured in high-glucose Dulbecco’s Modified Eagle Medium (DMEM; Thermo Fisher Scientific) supplemented with 10% fetal bovine serum (FBS; Sigma-Aldrich) and 1% penicillin–streptomycin (Gibco) at 37 °C in a humidified atmosphere containing 5% CO₂. Cells were routinely tested for mycoplasma contamination and used at passages below P15 to avoid replicative senescence.

For induction of stress-induced premature senescence (SIPS), MRC-5 cells were treated with doxorubicin (0.5 µM; Sigma-Aldrich) for 7 days. Where indicated, cells were treated with the p300/CBP inhibitor A485 (1 µM; Thermo Fisher Scientific), either alone or in combination with doxorubicin. Control cells received DMSO (0.1%).

Human umbilical vein endothelial cells (HUVECs) were isolated from anonymously acquired umbilical cords according to the ethical principles of the Declaration of Helsinki and with approval from the Jena University Hospital ethics committee (2023-2894-Material). Donors were informed and provided with written consent. HUVECs were isolated as previously described (Stabenow et al., 2022). Briefly, endothelial cells were detached from umbilical veins using collagenase (0.01% in M199, 3 min, 37 °C), suspended in M199 supplemented with 10% FCS, washed once by centrifugation (500 × g, 6 min), resuspended, and seeded onto culture flasks coated with 0.2% gelatine. After 24 h, medium was replaced with full growth medium consisting of M199 supplemented with 17.5% FCS, 2.5% human serum, 7.5 µg/mL ECGS, 7.5 U/mL heparin, 680 µM glutamine, 100 µM vitamin C, 100 U/mL penicillin, and 100 µg/mL streptomycin. Cells were cultured until confluence, detached using trypsin/EDTA (3 min, 37 °C), split, and reseeded. Replicative senescence was induced by serial passaging of primary cells. Half of each passage was frozen in liquid nitrogen. For experiments, primary cells and cells from passages 5 and 20 were thawed and grown to confluence before analysis (Stabenow et al., 2022).

#### Senescence and viability assays

Senescence-associated β-galactosidase (SA-β-gal) staining was performed using a pH 6.0 staining solution following fixation with 4% paraformaldehyde. Cells were incubated at 37 °C until signal development and quantified by manual counting of at least five random fields per condition. Cell density and viability were assessed by crystal violet staining (0.1%, 30 min, room temperature), followed by solubilization and absorbance measurement at 570 nm. All assays were performed with three independent biological replicates (n = 3).

#### Immunofluorescence microscopy

Cells were seeded on glass coverslips, fixed with paraformaldehyde (Roti-Histofix; Carl Roth), and permeabilized with 0.2% Triton X-100. After blocking, samples were incubated with primary antibodies against intracellular trafficking markers (EEA1, SEC31A, LAMP1, ERGIC-53, Giantin, TGN46), followed by fluorophore-conjugated secondary antibodies (Alexa Fluor series). Nuclei were stained with Hoechst 33342. Images were acquired using a Zeiss ApoTome optical sectioning microscope under identical acquisition settings across conditions. Experiments were performed with three independent biological replicates (n = 3).

#### Protein extraction and immunoblotting

Cells were lysed in STEN buffer supplemented with protease inhibitors, and protein concentration was determined using a BCA assay. Equal amounts of protein were separated by SDS-PAGE and transferred to PVDF membranes. Membranes were incubated with primary antibodies followed by HRP-conjugated secondary antibodies and visualized using enhanced chemiluminescence (ECL).

#### Proteomics and PTM analysis

Whole-cell proteomics and post-translational modification (PTM) analyses were performed at the Proteomics Facility of the Leibniz Institute on Aging (FLI, Jena). Sample preparation for PTMs and whole cells was performed according to Marino et al. (Marino et al., 2025) with slight changes (see detailed method description in the supplementary section). Whole cell samples were processed and analyzed using an Evosep One liquid chromatography system coupled to an Orbitrap Exploris 480 mass spectrometer (Thermo Fisher Scientific) operated in data-independent acquisition (DIA) mode. PTM samples were analyzed uning a NanoAcquity M-Class (Waters) coupled either to and Orbitrap Trihibryd Lumor or and Orbitrap Exploris 480 (Thermo Fisher Scientific). Acetylation, phosphorylation, and ubiquitination datasets were analyzed in parallel with the proteome. PTM intensities were normalized to corresponding protein abundance to distinguish modification-specific regulation from protein-level changes. Data were processed using Spectronaut (Biognosys) with a directDIA workflow against a human SwissProt database. False discovery rate (FDR) was controlled at 1% at both peptide and protein levels. Biological replicates included: n = 5 for DMSO, DOXO, and contact inhibition (CI) comparisons (MRC-5 and HUVEC datasets); n = 4 for experiments involving A485 treatment. A detailed method description can be found in the supplementary section.

#### Secretome analysis

For secretome profiling, cells were washed and incubated in serum-free medium for 4 h. Conditioned media were collected, supplemented with protease inhibitors, and cleared of debris by centrifugation. Secretome samples were normalized to cell number prior to analysis.

#### Bioinformatic and statistical analysis

Principal component analysis (PCA), differential expression analysis, and correlation analyses were performed using R. Gene set enrichment analysis (GSEA) and over-representation analysis (ORA) conducted using WebGestalt and Reactome pathway databases. Network analyses were performed using Cytoscape with the ClueGO plugin and STRING. Statistical analyses were performed using unpaired t-tests or one-way ANOVA as indicated in figure legends. Data are presented as mean ± standard deviation (SD).

## Supporting information

contains suppl. info and suppl. figures

## Acknowledgements

We thank the FLI Imaging Core Facility for support with microscopy and image acquisition and Fabian Monheim and Philipp Koch from FLÍs Core Facility Life Science Computing for their support in developing and implementing the interactive senescence PTM atlas. This work was supported by the Deutsche Forschungsgemeinschaft (DFG)-funded Research Training Group ProMoAge (RTG 2155) and by the Leibniz Graduate School on Aging (LGSA).

## Author Contributions

T.N. conceived the study, performed experiments, analyzed data, prepared figures, and wrote the original manuscript draft. B.E. generated and characterized cellular senescence samples used in comparative analyses. R.H. supervised these contributions and provided scientific input. E.C., N.R., and N.P. contributed proteome study design, analysis, and experimental troubleshooting. J.G. and C.G. contributed secretome study design, analysis, and interpretation. C.K. conceived and supervised the study, secured funding, interpreted data, and revised the manuscript. All authors reviewed and approved the final manuscript.

## Conflict of Interest

JG is co-founder, shareholder and scientific advisor of Rockfish Bio. All other authors declare no competing interests.

## Data Availability Statement

The mass spectrometry proteomics datasets generated during this study have been deposited in the MassIVE repository. Whole-cell proteome and PTM datasets: accession **MSV000101788, MSV000101789, MSV000101790, MSV000101791**. Secretome data have been deposited to the ProteomeXchange Consortium via the PRIDE (Perez-Riverol et al., 2025) partner repository with the dataset identifier PXD080515. Additional data supporting the findings of this study are available from the corresponding author upon reasonable request.

## Notes

https://genome.leibniz-fli.de/senescence-ptm-atlas/

### Summary of Updates

To make the large-scale datasets from this study more accessible, we developed an interactive senescence PTM atlas, which allows users to explore protein abundance and PTM changes across the senescence models and conditions analyzed in the manuscript. A link to that atlas was added at the end of the results section. In the previous version the suppl. info was missing, this was corrected.

