## Supplementary material for "Acetylation-dependent remodeling of the secretory pathway shapes the senescence-associated secretome": contains suppl. info and suppl. figures

[5] Austrian Cluster for Tissue Regeneration

Content:

1. Extended proteomics methods

2. Legends suppl. figures

3. Suppl. Figures 1-9

### Proteomics extended methods

#### 1. Sample Preparation for Proteome and Analysis of PTMs

To prepare samples for proteomic analysis, cells were lysed in a buffer comprised of 5% SDS, 100 mM HEPES, and 50 mM DTT. The resulting mixture was then subjected to sonication using Bioruptor Plus (Diagenode, Belgium) for a high-intensity cycle of 10 repeats, consisting of 30 seconds on and 60 seconds off at a maintained temperature of 20°C. This was promptly followed by a thermal denaturation step where samples were heated at 95°C for 5 minutes. Thiol groups in the proteins were reduced and then alkylated by adding iodoacetamide to a final concentration of 15 mM and incubated in darkness at room temperature for 30 minutes. After reduction and alkylation, samples were acidified with phosphoric acid to a 2.5% final concentration. The sample was mixed with the s-trap binding buffer containing 90% methanol and 100 mM Tetraethylammonium bromide (TEAB), totaling 1395 $\mu$L. The sample buffer mix was applied to a 96-well S-trap microplate (Protifi) to capture proteins, followed by three wash cycles with the binding buffer. Samples were then digested enzymatically with trypsin (1 $\mu$g per sample in 50 mM TEAB, pH 7.55) and incubated at 47°C for 1 hour. For peptide elution, a three-step procedure was employed using (1) 50 mM TEAB at pH 7.55, (2) elution buffer 1 with 0.2% formic acid in water, and (3) elution buffer 2 comprising 50% acetonitrile with 0.2% formic acid. Eluted peptides were concentrated and desalted via vacuum centrifugation using an Eppendorf Concentrator Plus (Eppendorf AG, Germany). The dried samples were reconstituted in Evosep buffer A (0.1% formic acid in water) and subjected to additional sonication on Bioruptor Plus for three cycles at 60 seconds on and 30 seconds off at 20°C. Following sonication, peptide samples were processed for Evotip-based LC-MS analysis. Evotips were prepared by washing with Evosep buffer B (acetonitrile with 0.1% formic acid), conditioning with isopropanol, and equilibrating with Evosep buffer A as per the manufacturer's instructions. Peptide samples were then loaded onto the prepared Evotips, washed to remove non-peptide material with Evosep buffer A, and the tips were filled with the same buffer in readiness for mass spectrometric analysis.

For PTM enrichment, peptides were further processed as described below. For Data Independent Acquisition (DIA) based analysis of total proteome, samples were transferred to MS vials, diluted to a concentration of 1 µg/µL, and spiked with iRT kit peptides (Biognosys, Ki-3002-2) before analysis by LC-MS/MS.

#### 2. Sequential Enrichment of Ubiquitylated and Acetylated Peptides

Ubiquitylated and acetylated peptides were sequentially enriched starting from ~1000 µg of dried peptides per replicate. For the enrichment of ubiquitylated peptides, the PTMScan® HS Ubiquitin/SUMO Remnant Motif (K-ε-GG) kit (Cell Signaling Technology, 59322) was used following manufacturer instructions. The K-ε-GG modified enriched fraction was desalted and concentrated as described above, dissolved in MS buffer A, and spiked with iRT kit peptides before LC-MS/MS analysis.

The flowthrough fractions from the K- ε -GG enrichment were acidified with 10% (v/v) trifluoroacetic acid and desalted using Oasis® HLB µElution Plate 30 µm (30 mg) following manufacturer instructions. Acetylated peptides were enriched as described by (Di Sanzo et al. 2021). Briefly, dried peptides were dissolved in 1000 µl of IP buffer (50 mM MOPS pH 7.3, 10 mM KPO4 pH 7.5, 50 mM NaCl, 2.5 mM Octyl β-D-glucopyranoside) to reach a peptide concentration of 1 µg/µL, followed by sonication in a Bioruptor Plus (5 cycles with 1 min ON and 30 s OFF with high intensity at 20 °C). Agarose beads coupled to an antibody against acetyl-lysine (ImmuneChem Pharmaceuticals Inc., ICP0388-5MG) were washed three times with washing buffer (20 mM MOPS pH 7.4, 10 mM KPO4 pH 7.5, 50 mM NaCl) before incubation with each peptide sample for 1.5 h on a rotating well at 750 rpm (STARLAB Tube 19 roller Mixer RM Multi-1). Samples were transferred into Clearspin filter microtubes (0.22 µm) (Dominique Dutscher SAS, Brumath, 007857ACL) and centrifuged at 4 °C for 1 min at 2000 x*g*. Beads were washed first with IP buffer (three times), then with washing buffer (three times), and finally with 5 mM ammonium bicarbonate (three times). Thereupon, the enriched peptides were eluted first in basic condition using 50 mM aqueous NH3, then using 0.1% (v/v) trifluoroacetic acid in 10% (v/v) 2-propanol and finally with 0.1% (v/v) trifluoroacetic acid. Elutions were dried down and reconstituted in MS buffer A (5% (v/v) acetonitrile, 0.1% (v/v) formic acid), acidified with 10% (v/v) trifluoroacetic acid, and then desalted with Oasis® HLB µElution Plate 30 µm. Desalted peptides were finally dissolved in MS buffer A, spiked with iRT kit peptides, and analyzed by LC-MS/MS.

#### 3. Enrichment of Phosphorylated Peptides

Desalted peptides corresponding to 200 µg, as described in "Sample preparation for total proteome and analysis of PTMs' were used. The last desalting step was performed using 50 µl of 80% ACN and 0.1% TFA buffer solution. Before phosphopeptide enrichment, samples were filled up to 210 µl using 80% ACN and 0.1% TFA buffer solution. Phosphorylated peptides were enriched using Fe(III)-NTA cartridges (Agilent Technologies, G5496-60085) in an automated fashion using the standard protocol from the AssayMAP Bravo Platform (Agilent Technologies). In short, Fe(III)-NTA cartridges were first primed with 100 µl of priming buffer (100% ACN, 0.1% TFA) and equilibrated with 50 µL of buffer solution (80% ACN, 0.1% TFA). After loading the samples into the cartridge, the cartridges were washed with an OASIS elution buffer, while the syringes were washed with a priming buffer (100% ACN, 0.1% TFA). The phosphopeptides were eluted directly with 25 µL of 1% ammonia into 25 µL of 10% FA. Samples were dried with a speed vacuum centrifuge and stored at –20 °C until LC-MS/MS analysis.

#### 4. LC-MS Data Independent Analysis (DIA)

For long gradient acquisition, ubiquitin analysis and acetylation analysis, Approximatively 1µg of reconstituted were separated using a nanoAcquity UPLC (Waters, Milford, MA) was coupled online to the MS. Peptide mixtures were separated in trap/elute mode, using a trapping (nanoAcquity Symmetry C18, 5 µm, 180 µm × 20 mm) and an analytical column (nanoAcquity BEH C18, 1.7 µm, 75 µm x 250 mm). The outlet of the analytical column was coupled directly to an Orbitrap Fusion Lumos mass spectrometers (Thermo Fisher Scientific, San Jose, CA) using the Proxeon nanospray source. Solvent A was water, 0.1% formic acid and solvent B was acetonitrile, 0.1% formic acid. The samples were loaded with a constant flow of solvent A, at 5 µL/min onto the trapping column. Trapping time was 6 min. Peptides were eluted via the analytical column with a constant flow of 300 nL/min. During the elution step, the percentage of solvent B increased in a nonlinear fashion from 0% to 40% in 120 min. Total runtime was 145 min, including cleanup and column re-equilibration. The peptides were introduced into the mass spectrometer via a Pico-Tip Emitter 360 $\mu$m OD x 20 $\mu$m ID; 10 $\mu$m tip (New Objective) and a spray voltage of 2.2 kV was applied. The capillary temperature was set at 300 °C. The RF lens was set to 30%.

Full scan MS spectra with mass range 350-1650 m/z were acquired in profile mode in the Orbitrap with resolution of 120,000 FWHM. The filling time was set at maximum of 20 ms with an AGC target of 5 x $10^5$ ions. DIA scans were acquired with 40 mass window segments of differing widths across the MS1 mass range. The HCD collision energy was set to 30%. MS/MS scan resolution in the Orbitrap was set to 30,000 FWHM with a fixed first mass of 200m/z after accumulation of 1x $10^6$ ions or after filling time of 70ms (whichever occurred first). Data were acquired in profile mode. For data acquisition and processing Tune version 2.1 and Xcalibur 4.1 were employed.

For short gradient avquisition, Peptide separation and analysis were accomplished by the Proteomics facility at the FLI using the Evosep One chromatography system (Evosep, Odense, Denmark), which was outfitted with an Evosep Endurance column, 15 cm x 150 µm internal diameter packed with 1.9 µm Reprosil-Pur C18 beads (EV-1106, PepSep, Marslev, Denmark). Chromatographic separation was achieved using a specific 44-minute gradient designed for a throughput of 30 samples per day; this proprietary gradient utilized solvent A (water with 0.1% formic acid) and solvent B (acetonitrile with 0.1% formic acid). Mass spectrometric analysis was conducted on an Orbitrap Exploris 480 instrument (Thermo Fisher Scientific, Bremen, Germany). This system was fitted with PepSep Sprayers and a Proxeon nanospray source for ionization. The sprayed peptides were introduced via a heated PepSep Emitter (360-µm outer diameter and 20-µm inner diameter) at 300°C with a 2.2 kV spray voltage. The instrument's injection capillary was also heated to a temperature of 300°C. The radio frequency of the ion funnel was set at 30% to focus the ions for improved detection. Data-independent acquisition (DIA) was implemented to capture comprehensive mass spectrometry (MS) spectra, ranging from 350 to 1650 m/z, with a high resolution of 120,000 FWHM in the Orbitrap detector. The system was programmed to handle a default charge state of 2+ and to enforce a maximum injection time of 60 ms or an ion target of 3 × 10^6, whichever was reached first. The DIA MS/MS method involved segmenting the mass range into 40 windows of varying widths. Fragmentation was achieved using higher collisional dissociation with stepped normalized collision energies (NCE) of 25%, 27.5%, and 30%. The resulting MS/MS spectra were acquired at a resolution of 30,000 FWHM, ensuring a minimum m/z of 200. Raw data were collected in profile mode and processed using Xcalibur software version 4.4 and Tune version 3.1 (Thermo Fisher Scientific).

For phosphopeptides, samples were reconstituted in in MS Buffer (5% acetonitrile, 95% Milli-Q water, with 0.1% formic acid) and spiked with iRT peptides (Biognosys, Switzerland). Peptides were separated in trap/elute mode using the nanoAcquity MClass Ultra-High Performance Liquid Chromatography system (Waters, Waters Corporation, Milford, MA,USA ) equipped with a trapping (Waters nanoEase M/Z Symmetry C18, 5µm, 180 µm x 20 mm) and an analytical column (Waters nanoEase M/Z Peptide C18, 1.7µm, 75µm x 250mm). Solvent A was water and 0.1% formic acid, and solvent B was acetonitrile and 0.1% formic acid. 5 $\mu$l of the sample were loaded with a constant flow of solvent A at 5 µl/min onto the trapping column. Trapping time was 6 min. Peptides were eluted via the analytical column with a constant flow of 0.3 µl/min. During the elution step, the percentage of solvent B increased in a nonlinear fashion from 0–40% in 60 min. Total run time was 75 min.
The LC was coupled to an Orbitrap Exploris 480 (Thermo Fisher Scientific, Bremen, Germany) using the Proxeon nanospray source. The peptides were introduced into the mass spectrometer via a Pico-Tip Emitter 360-µm outer diameter × 20-µm inner diameter, 10-µm tip (New Objective) heated at 300 °C, and a spray voltage of 2.2 kV was applied. The capillary temperature was set

at 300°C. The radio frequency ion funnel was set to 30%. Full scan mass spectrometry (MS) spectra with mass range 350–1650 m/z were acquired in profile mode in the Orbitrap with resolution of 120,000 FWHM. The default charge state was set to 3+. The filling time was set at maximum of 60 ms with limitation of $3 \times 10^6$ ions. DIA scans were acquired with 40 mass window segments of differing widths across the MS1 mass range. Higher collisional dissociation fragmentation (stepped normalized collision energy; 25, 27.5, and 30%) was applied and MS/MS spectra were acquired with a resolution of 30,000 FWHM with a fixed first mass of 200 m/z after accumulation of $3 \times 10^6$ ions or after filling time of 35 ms (whichever occurred first). Data were acquired in profile mode. For data acquisition and processing of the raw data Xcalibur 4.4 (Thermo) and Tune version 3.1 were used.

**5. Processing of Proteomic Data**

Raw data obtained from data-independent acquisition (DIA) were processed by the Proteomics facility at the FLI using the directDIA workflow in Spectronaut software (version 18, Biognosys, Switzerland). The datasets were matched against a species-specific SwissProt database tailored for Homo sapiens, comprising 20,816 protein entries, as well as a contaminant database with 247 entries. Protein identification considered variable modifications such as methionine oxidation (M) and N-terminal protein acetylation (acetyl). A tolerance for up to two missed cleavages by trypsin and a maximum of five variable modifications per peptide were allowed. Confidence in protein identification was solidified by imposing a false discovery rate (FDR) of 1% at both peptide and protein levels using target-decoy approaches. The quantitative analysis within Spectronaut employed the label-free quantification (LFQ) QUANT 2.0 method, incorporating Global Normalization. Precursor ions for quantification were filtered based on a 0.2 percentile fraction, and missing values were addressed with global imputation to ensure comprehensive relative abundance assessments. Further relative quantification was conducted pairwise between replicate sample sets from the various experimental conditions. Resulting data tables, alongside protein quantity reports, were exported for downstream analysis. Data analyses, including the selection of statistically significant protein changes, were conducted in R studio using customized pipelines and scripts developed in-house. Significant protein abundance changes were determined using a cutoff of a log2 fold change (log2FC) of 0.58 and a q-value threshold of ≤0.05 for all the experiments including PTM enrichment except for the HUVEC PTM enrichment data set where we had to use p-value of ≤0.05.

**6. PTM Analysis Based on Processed Proteomic Data**

Global proteomic and PTM-enriched datasets were obtained from MRC-5 fibroblasts and HUVECs, both in control and senescent conditions. For each cell line, two complementary datasets were generated: a whole-cell proteome and a PTM-enriched proteome, including acetylation, phosphorylation, and ubiquitylation. PTM-specific intensities were normalized to the corresponding protein abundance derived from the whole-cell dataset to correct for changes in baseline protein expression.

PTM-specific analyses were then performed to determine the number of significantly modified proteins in senescent versus non-senescent cells for each modification type. For every PTM, the list of modified proteins was filtered for subcellular localization and GO term enrichment focused specifically on ER- and Golgi-associated processes and components, to selectively extract modification patterns in secretory pathway proteins.

To investigate potential competitive regulation between lysine acetylation and ubiquitylation, modified protein lists were cross-referenced to identify overlapping modification sites within the same protein substrates. In parallel, the distribution of $\log_2$ fold changes were analyzed across all PTM datasets to assess the relative prevalence of up- or downregulation of each modification type under senescent conditions.

Comparative analysis between MRC-5 and HUVEC datasets emphasized acetylation, which was the most prominently regulated PTM in both models. Shared acetylation events on ER- and Golgi-localized proteins were identified to determine common PTM signatures associated with senescence in the secretory pathway.

To guide the selection of a candidate protein for validation, biological process enrichment was performed using Cytoscape with the ClueGO plugin, applied both to the global PTM-regulated proteome and specifically to the subset of ER/Golgi-localized proteins. Resulting GO terms and associated protein lists were compiled to identify pathways and molecular functions of interest. Based on the overlap in modification sites, functional enrichment, and cross-cell line consistency, a target protein was selected and evaluated using a Parallel Reaction Monitoring (PRM) approach to validate the significance and regulation of its PTM sites.

All graphs and data visualizations derived from proteomics and PTM datasets were generated either using R (version 4.4.2) within the RStudio environment, employing standard packages for data handling and graphical representation or using combination of Microsoft Excel and GraphPad Prism (version 10).

### Supplementary figure legends

Supplementary Figure 1. Immunofluorescence analysis of secretory pathway markers in MRC-5 cells following doxorubicin treatment.

MRC-5 fibroblasts were treated with vehicle control (DMSO, 0.1%) or doxorubicin (DOXO, 0.5 $\mu$M) for 7 days, fixed, and immunostained for ERGIC-53 (ER–Golgi intermediate compartment), GIANTIN (Golgi), or TGN46 (trans-Golgi network). Secondary antibodies conjugated to Alexa Fluor 488 were used for ERGIC-53 and GIANTIN, whereas TGN46 was detected using Alexa Fluor 555. Images represent single optical sections acquired under identical imaging conditions using a Zeiss Apotome optical sectioning system. Representative images are shown.

Supplementary Figure 2. Replicative Senescence Reprograms the HUVEC Whole Cell and PTM Proteome Across Passages.

(A) Principal component analysis (PCA) of whole-cell proteomics comparing early (P1), intermediate (P5), and late (P20) passage HUVECs.
(B) Principal component analysis (PCA) of acetylation and phosphorylation datasets comparing early (P1), intermediate (P5), and late (P20) passage HUVECs (n = 5 per condition).
(C) Total number of significantly regulated proteins ($log_2FC \leq -0.58$ or $\geq 0.58$, Qvalue $\leq 0.05$) between condition comparisons.
(D) Number of identified features (significant and non-significant protein signals detected by MS) in each PTM category and across the whole proteome.
(E) Scatterplot describing the correlation between proteome and acetylome, in orange are highlighted proteins that are affected significantly in both layers, in purple proteins which are significantly changing only for acetylation.
(F) Bar chart showing distribution of significantly modified proteins into two subsets, ER and Golgi proteins in orange, Non-ER and Golgi in purple.
(G) Distribution of up- and downregulated modification sites across PTMs in the whole proteome.
(I) Distribution of up- and downregulated modification sites across PTMs in ER- and Golgi-associated proteins.

Supplementary Figure 3. Senescence-Associated PTM Sites Are Conserved Between MRC-5 and HUVEC Models.

(A) Comparison of chemically induced senescent MRC-5 (DMSO vs DOXO) and replicatively induced senescent HUVEC (P1 vs P20) datasets showing the number of acetylated proteins modified at identical sites across both models.
(B) Same-site acetylation overlap restricted to ER- and Golgi-associated proteins.
(C) Comparison of chemically induced senescent MRC-5 (DMSO vs DOXO) and replicatively induced senescent HUVEC (P1 vs P20) datasets showing the number of phosphorylated proteins modified at identical sites across both models.
(D) Same-site phosphorylation overlap restricted to ER- and Golgi-associated proteins.

Supplementary Figure 4. GO enrichment of proteins sharing identical acetylation or phosphorylation sites across MRC-5 and HUVEC senescence models.

(A) Enrichment results for proteins sharing identical acetylation sites across both cell lines. (B) Enrichment results for proteins sharing identical phosphorylation sites across both cell lines. Bubble size reflects the number of genes contributing to each term, and color indicates FDR.

Supplementary Figure 5. Doxorubicin Activates p300/CBP and Elevates Histone H3 Acetylation.

(A) Schematic of human p300 showing major domains and the HAT domain with the autoacetylation loop (~aa 1520–1560) highlighted; lysins K1542 and K1546 (within the loop) that are significantly acetylated in DOXO vs DMSO are indicated in green.
(B) Quantification of acetylation levels at three lysine residues (K1542, K1546, K1590) on p300, identified from acetylation enrichment proteomics of DOXO vs DMSO compared to protein levels of p300 identified from whole cell proteomics of DOXO vs DMSO. Intensities were normalized to DMSO controls.
(C) Quantification of acetylation levels at five lysine residues (K9, K14, K18, K27, K23) on H3, identified from acetylation enrichment proteomics of DOXO vs DMSO compared to protein levels of H3 identified from whole cell proteomics of DOXO vs DMSO. Intensities were normalized to DMSO controls.

Panels B-C, n=5. Statistical significance was assessed using unpaired t test. Significance indicators: ****$p \leq 0.0001$; ***$p \leq 0.001$; **$p \leq 0.01$; *$p \leq 0.05$; ns, not significant.

Supplementary Figure 6. p300/CBP Inhibition Differentially Rewires Proliferation and Apoptosis Related Pathways.

(A) MS-based quantification of CDK4 protein levels across four conditions (DMSO, DOXO, A485, A485+DOXO). Intensities were normalized to the mean of the DMSO control.
(B) MS-based quantification of CDK6 protein levels across the four conditions, plotted as normalized protein intensity.
(C) MS-based quantification of CASP3 abundance across conditions (DMSO, DOXO, A485, A485+DOXO).
(D) MS-based quantification of BAX levels across conditions, shown as normalized protein intensity.
(E) Lysates from MRC-5 fibroblasts treated with DOXO (0.5 $\mu$M) and/or A485 (1 $\mu$M) were subjected to SDS–PAGE and Western blot. Blots were probed with an antibody against full-length PARP; stain-free total protein obtained by 10% TCA precipitation served as the loading control.

Panels A-B n=4, panel E, n=3. The graphs show the mean + standard deviation (SD).

Statistical significance was assessed using one-way ANOVA. Significance indicators: ****$p \leq 0.0001$; ***$p \leq 0.001$; **$p \leq 0.01$; *$p \leq 0.05$; ns, not significant.

Supplementary Figure 7. A485 and Doxorubicin Differentially Modulate KAT/HDAC Levels and Global Lysine Acetylation.

(A) Heatmap showing differential expressions of acetyltransferases and deacetylases across all treatment conditions (DMSO, A485, DOXO, A485+DOXO), generated from proteomics data. Visualization created with BioRender. Numbers indicate log2fold.
(B) Lysates from the four conditions were subjected to SDS–PAGE and Western blot. Blots were probed with an anti–acetyl-lysine antibody (Acetyl-K); stain-free total protein detection with 10% TCA served as the loading control. Two prominent bands (~below and above 15 kDa) were quantified.
(C) Quantification of two visible bands of acetylated lysine expression below and above 15kD marker normalized to DMSO control.
Panels B–C, n = 5; panel A, n = 4. Bar graphs show densitometric quantification normalized to TCA staining. Statistical significance was assessed using one-way ANOVA. Significance indicators: ****$p \leq 0.0001$; ***$p \leq 0.001$; **$p \leq 0.01$; *$p \leq 0.05$; ns, not significant.

Supplementary Figure 8. A485 Reverses an ER/Golgi ER-Stress Protein Network in Senescent MRC-5 Cells.
Proteins differentially modulated by A485 in the presence of doxorubicin (DOXO) were analyzed in STRING to identify enriched biological processes.
(A) Gene ontology analysis of biological processes for proteins downregulated in the A485+DOXO condition compared to DOXO alone.
(B) Gene ontology analysis of biological processes for proteins upregulated in the A485+DOXO condition compared to DOXO alone.

Supplementary Figure 9. p300/CBP Inhibition Modulates Proteostasis and Secretory Machinery Associated to SASP.

(A) Principal component analysis (PCA) of whole-cell proteomics matched to secretome showing distribution and separation of the four conditions (DMSO, A485, DOXO, A485+DOXO).
(B) MS-based quantification of chaperone levels across the four conditions, plotted as normalized protein intensity.
(C) MS-based quantification of SASP associated secretory machinery protein levels across the four conditions, plotted as normalized protein intensity.
(D) Gene ontology analysis of biological processes for secreted proteins downregulated in the A485+DOXO condition compared to A485 alone.
Panels A-C n=4. The graphs show the mean + standard deviation (SD).
Statistical significance was assessed using one-way ANOVA. Significance indicators: ****$p \leq 0.0001$; ***$p \leq 0.001$; **$p \leq 0.01$; *$p \leq 0.05$; ns, not significant.

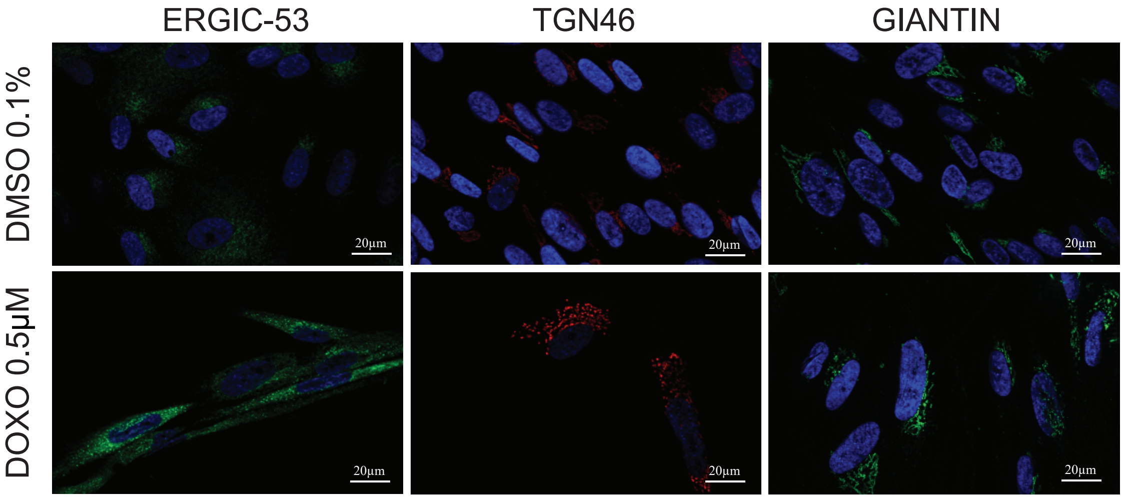


Suppl. Fig. 1

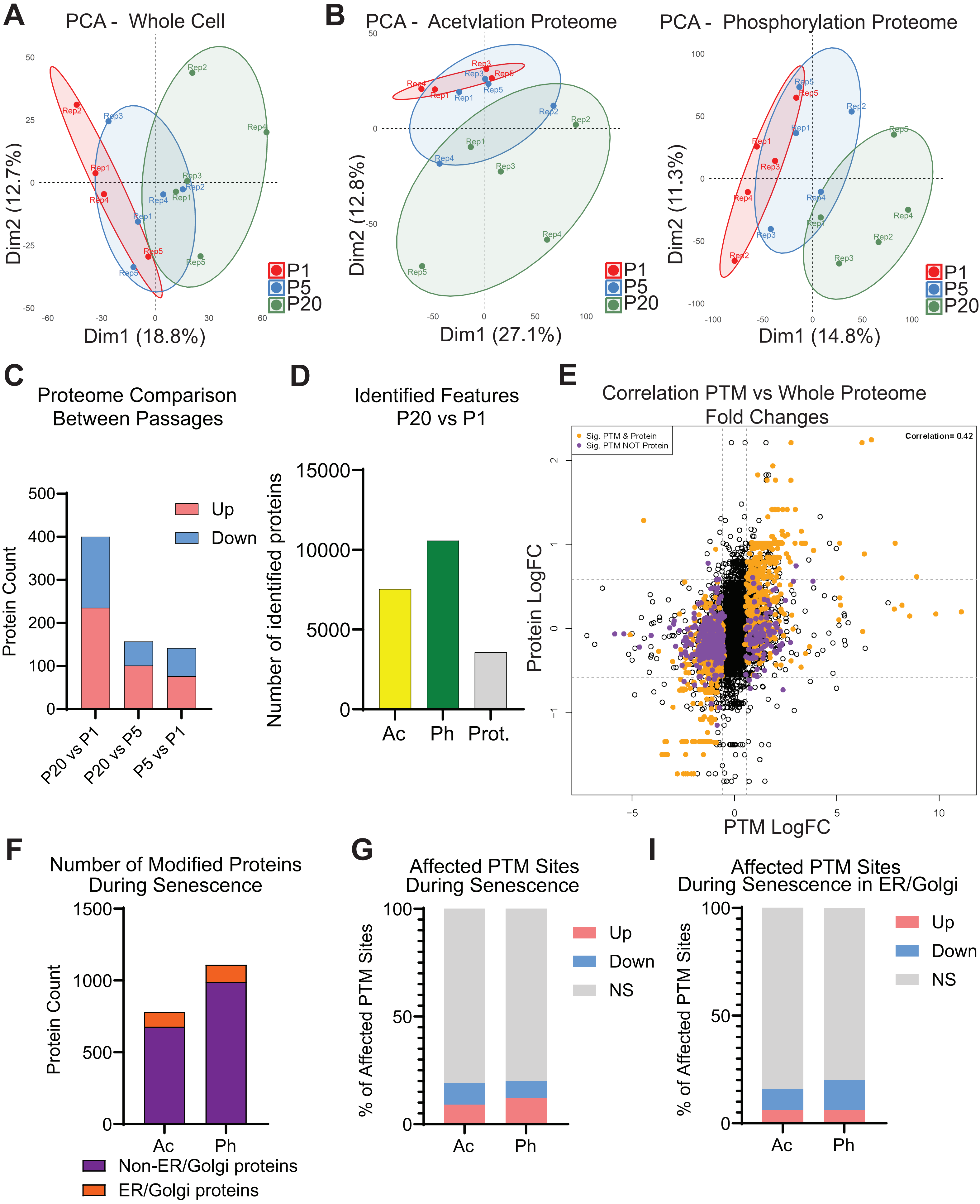
A
PCA - Whole Cell
Dim1 (18.8%)
Dim2 (12.7%)
P1
P5
P20
B
PCA - Acetylation Proteome
Dim1 (27.1%)
Dim2 (12.8%)
P1
P5
P20
PCA - Phosphorylation Proteome
Dim1 (14.8%)
Dim2 (11.3%)
P1
P5
P20
C
Proteome Comparison
Between Passages
Protein Count
Up
Down
P20 vs P1
P20 vs P5
P5 vs P1
D
Identified Features
P20 vs P1
Number of identified proteins
Ac
Ph
Prot.
E
Correlation PTM vs Whole Proteome
Fold Changes
Sig. PTM & Protein
Sig. PTM NOT Protein
Correlation= 0.42
Protein LogFC
PTM LogFC
F
Number of Modified Proteins
During Senescence
Protein Count
Ac
Ph
Non-ER/Golgi proteins
ER/Golgi proteins
G
Affected PTM Sites
During Senescence
% of Affected PTM Sites
Up
Down
NS
Ac
Ph
I
Affected PTM Sites
During Senescence in ER/Golgi
% of Affected PTM Sites
Up
Down
NS
Ac
Ph

Suppl. Fig. 2

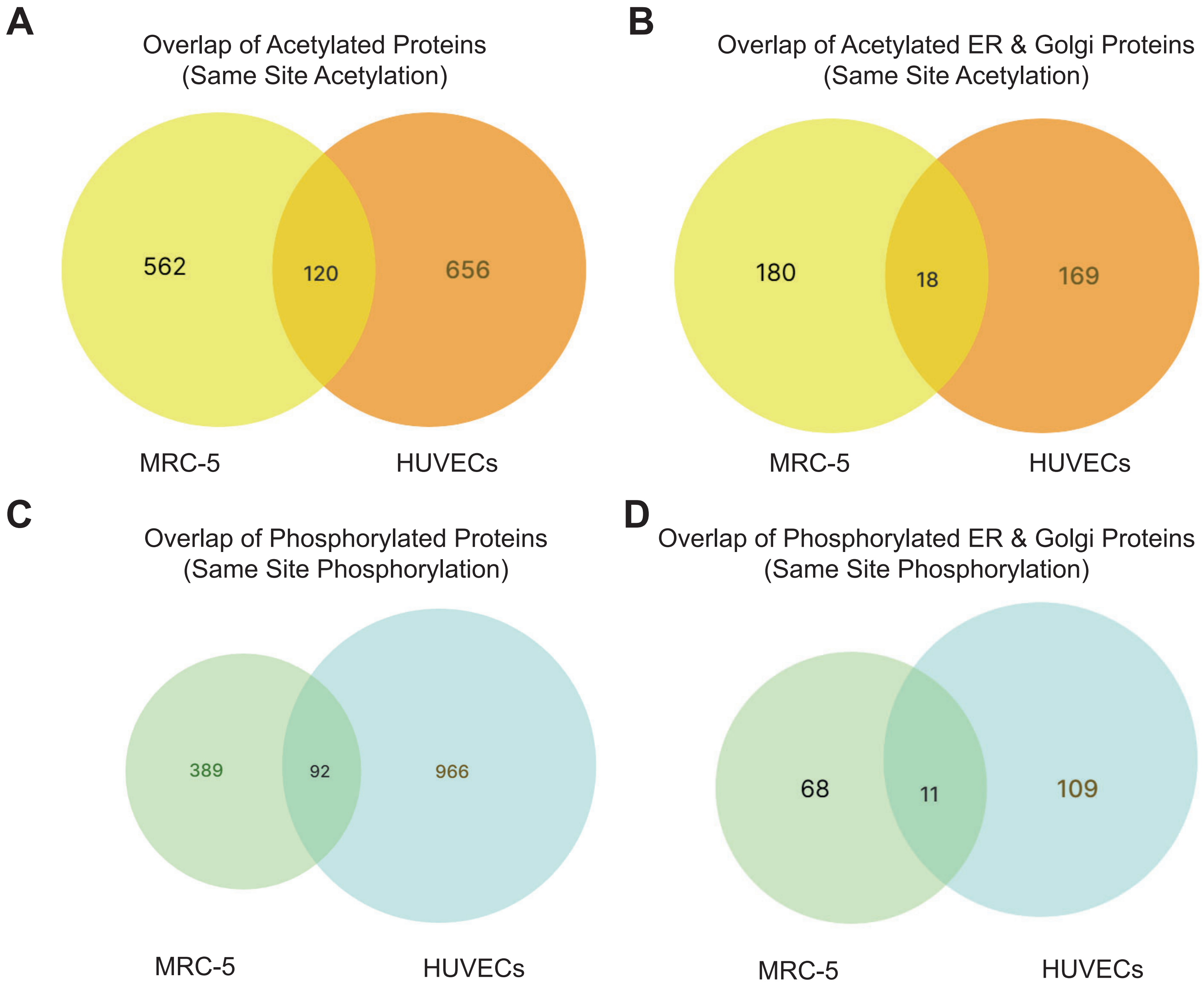
A
Overlap of Acetylated Proteins
(Same Site Acetylation)
562
120
656
MRC-5
HUVECs
B
Overlap of Acetylated ER & Golgi Proteins
(Same Site Acetylation)
180
18
169
MRC-5
HUVECs
C
Overlap of Phosphorylated Proteins
(Same Site Phosphorylation)
389
92
966
MRC-5
HUVECs
D
Overlap of Phosphorylated ER & Golgi Proteins
(Same Site Phosphorylation)
68
11
109
MRC-5
HUVECs

Suppl. Fig. 3

A

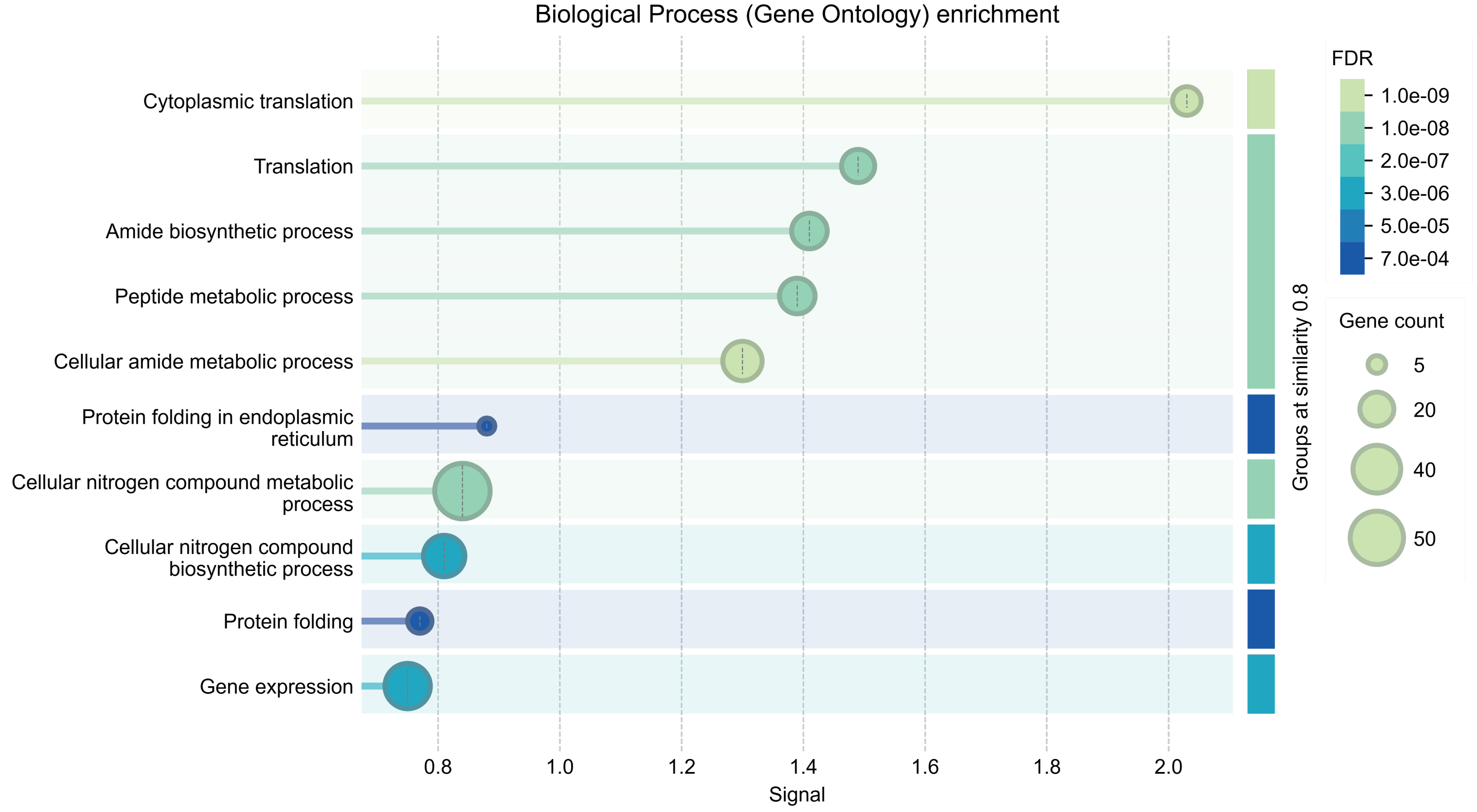


B

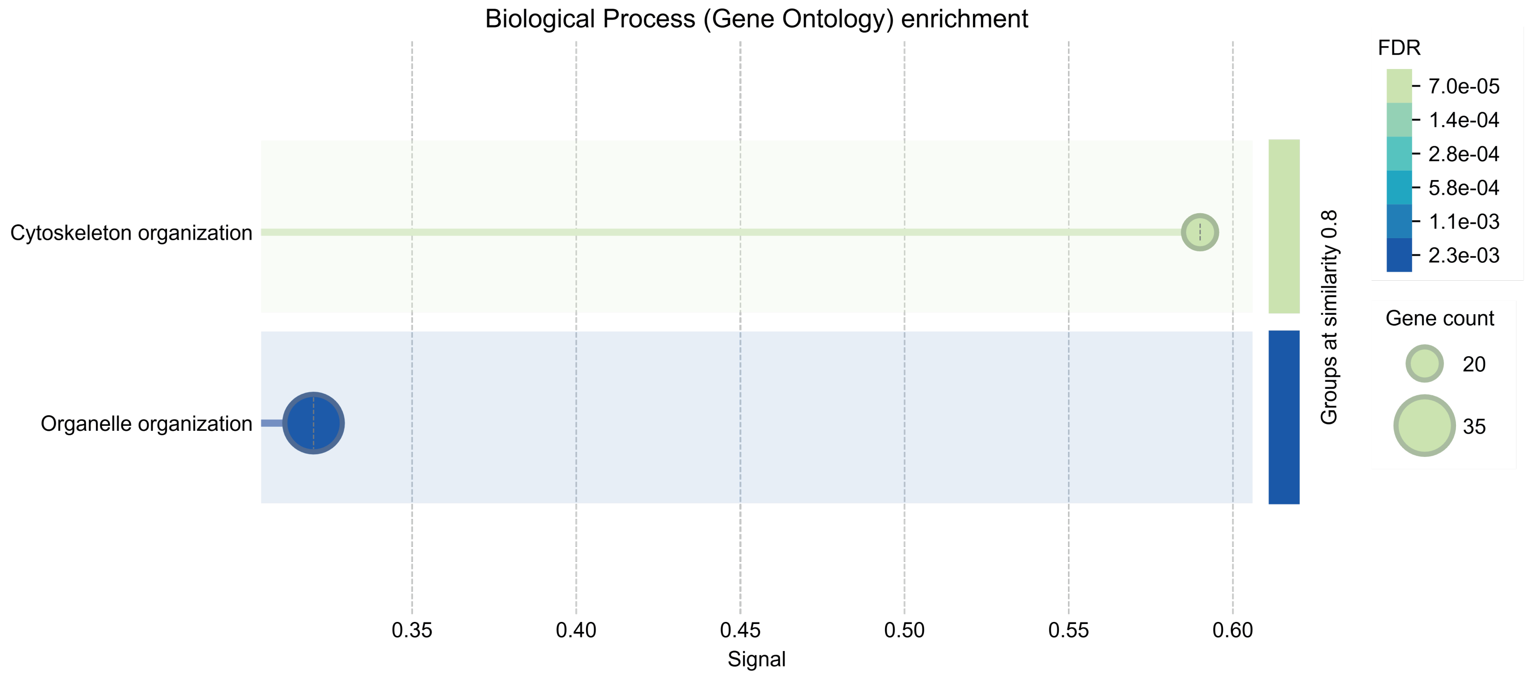


Suppl. Fig. 4

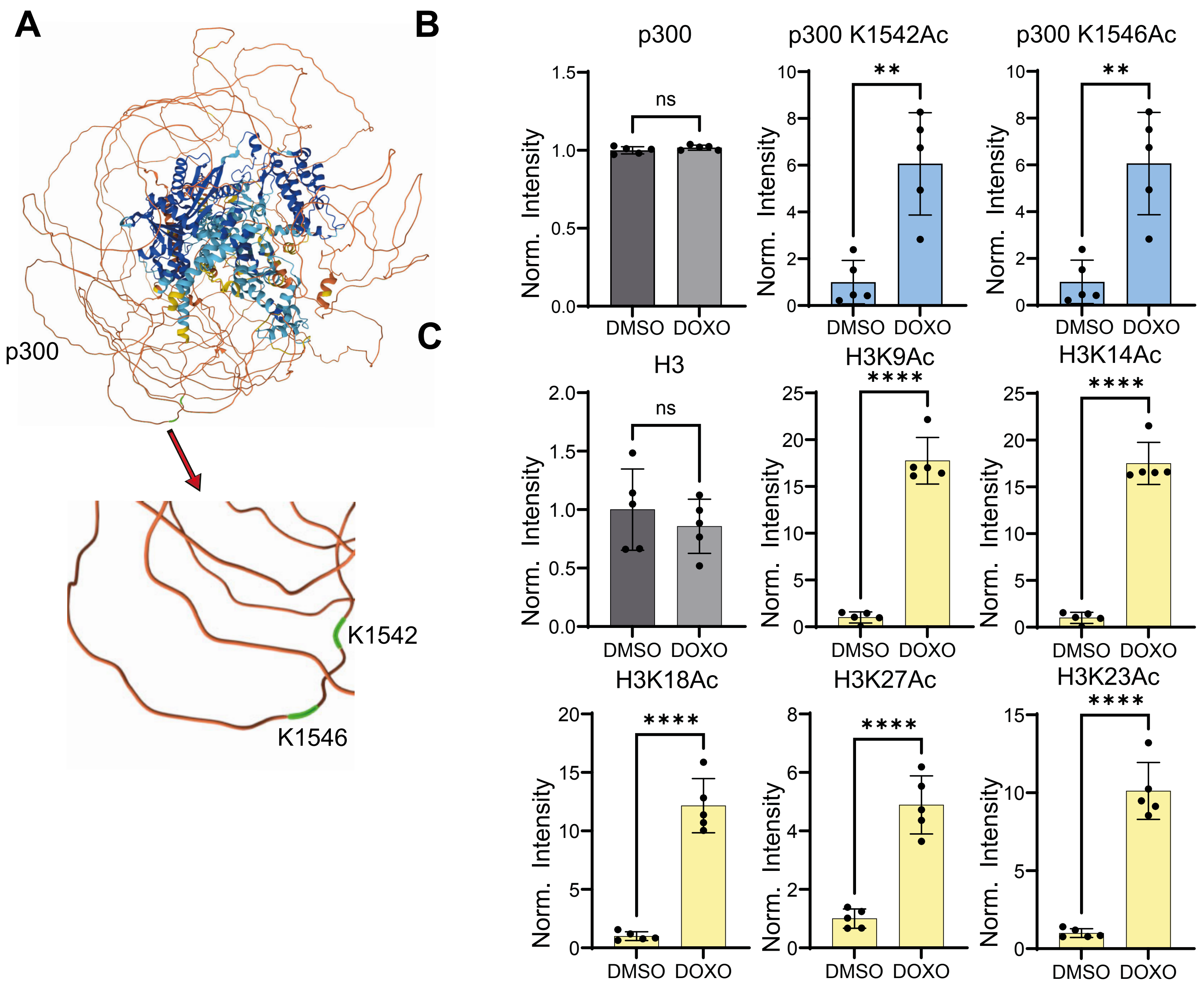
A
p300
K1542
K1546
B
p300
p300 K1542Ac
p300 K1546Ac
C
H3
H3K9Ac
H3K14Ac
H3K18Ac
H3K27Ac
H3K23Ac
Norm. Intensity
ns
**
****
DMSO
DOXO

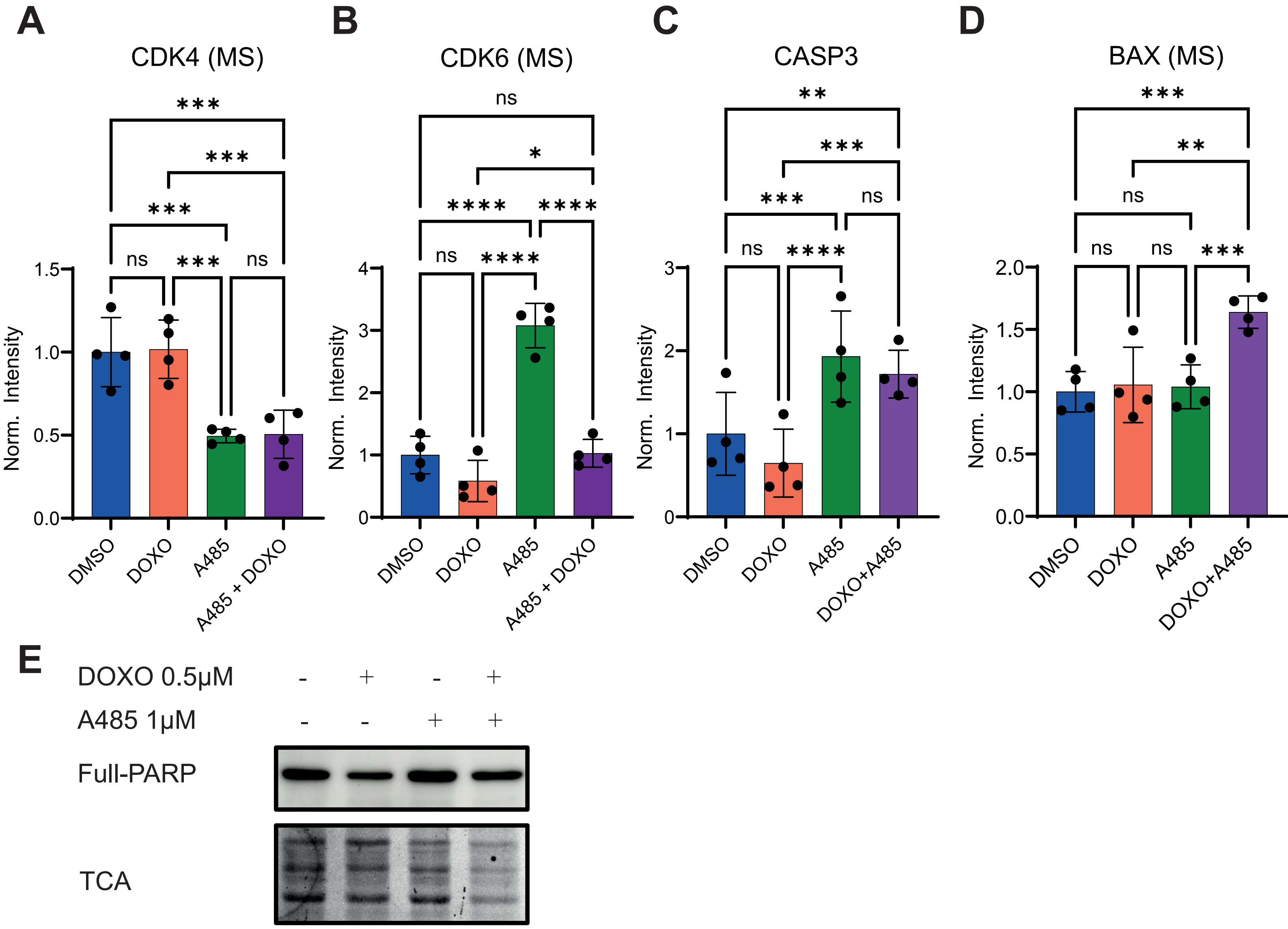
A
CDK4 (MS)
B
CDK6 (MS)
C
CASP3
D
BAX (MS)
Norm. Intensity
DMSO
DOXO
A485
A485 + DOXO
DOXO+A485
ns
*
**
***
****
E
DOXO 0.5μM
A485 1μM
Full-PARP
TCA

Suppl. Fig. 6

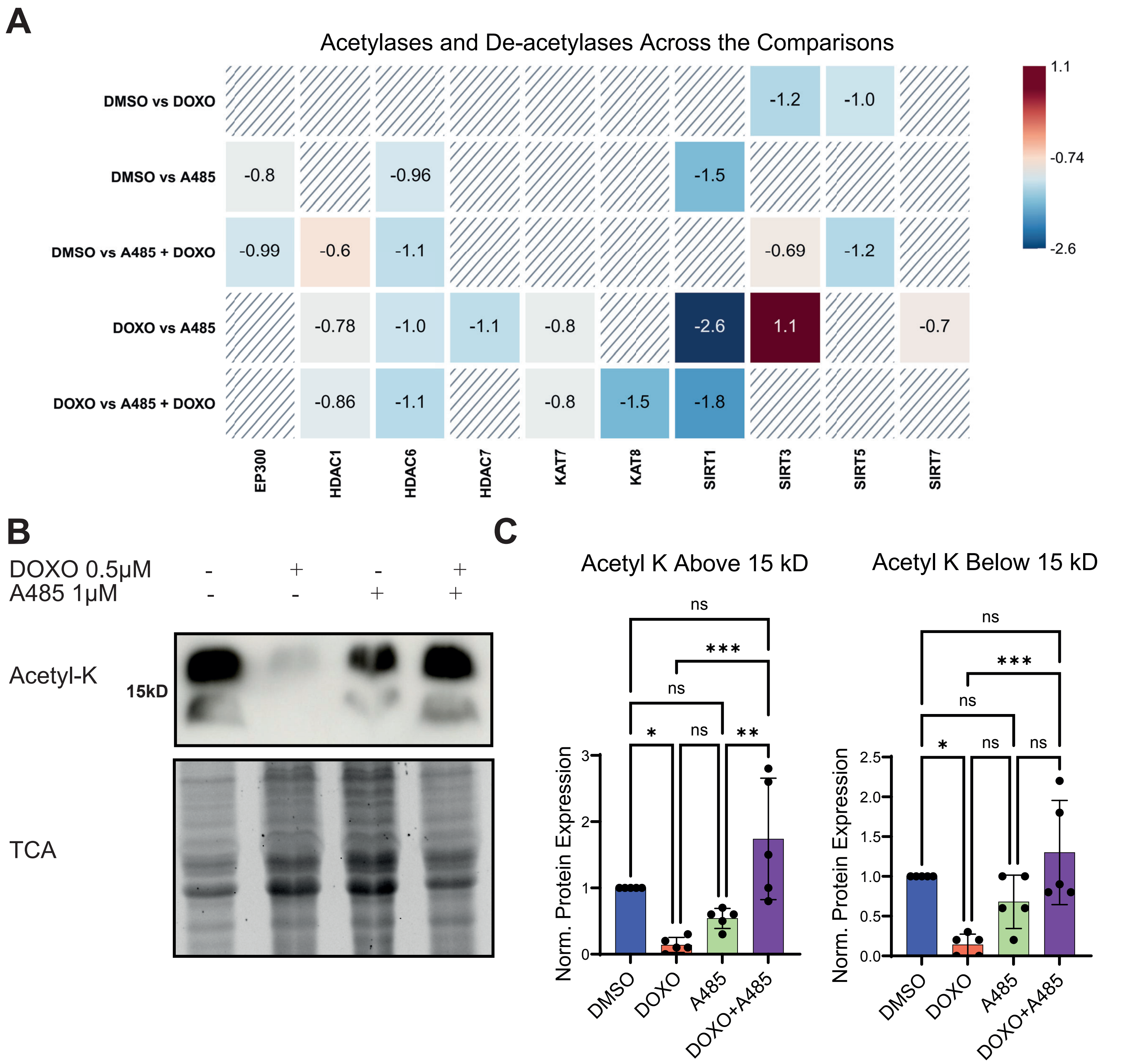
A
Acetylases and De-acetylases Across the Comparisons
DMSO vs DOXO
DMSO vs A485
DMSO vs A485 + DOXO
DOXO vs A485
DOXO vs A485 + DOXO
-1.2
-1.0
-0.8
-0.96
-1.5
-0.99
-0.6
-1.1
-0.69
-1.2
-0.78
-1.0
-1.1
-0.8
-2.6
1.1
-0.7
-0.86
-1.1
-0.8
-1.5
-1.8
EP300
HDAC1
HDAC6
HDAC7
KAT7
KAT8
SIRT1
SIRT3
SIRT5
SIRT7
1.1
-0.74
-2.6
B
DOXO 0.5μM - + - +
A485 1μM - - + +
Acetyl-K
15kD
TCA
C
Acetyl K Above 15 kD
Acetyl K Below 15 kD
ns
***
ns
*
ns
**
Norm. Protein Expression
DMSO
DOXO
A485
DOXO+A485

A

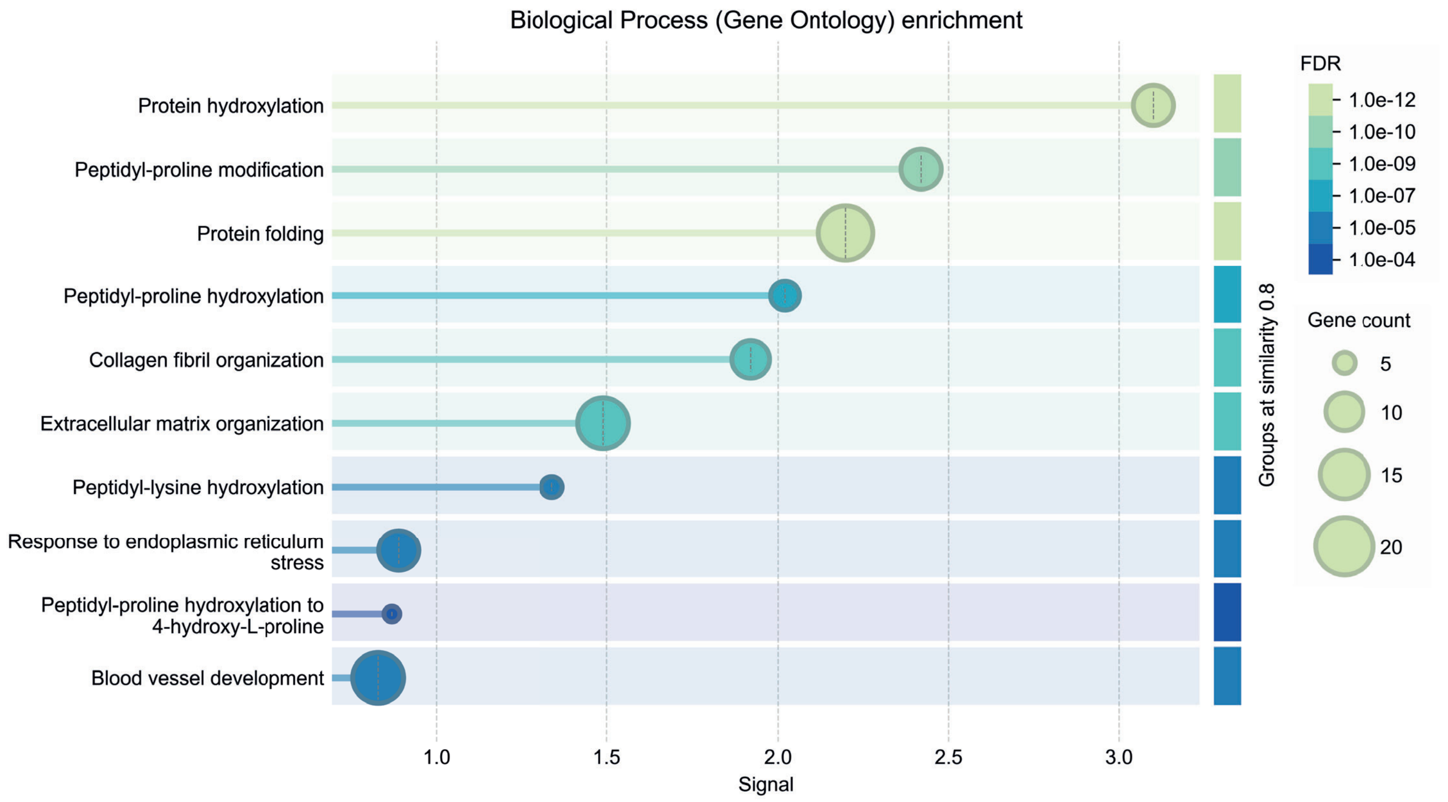


B

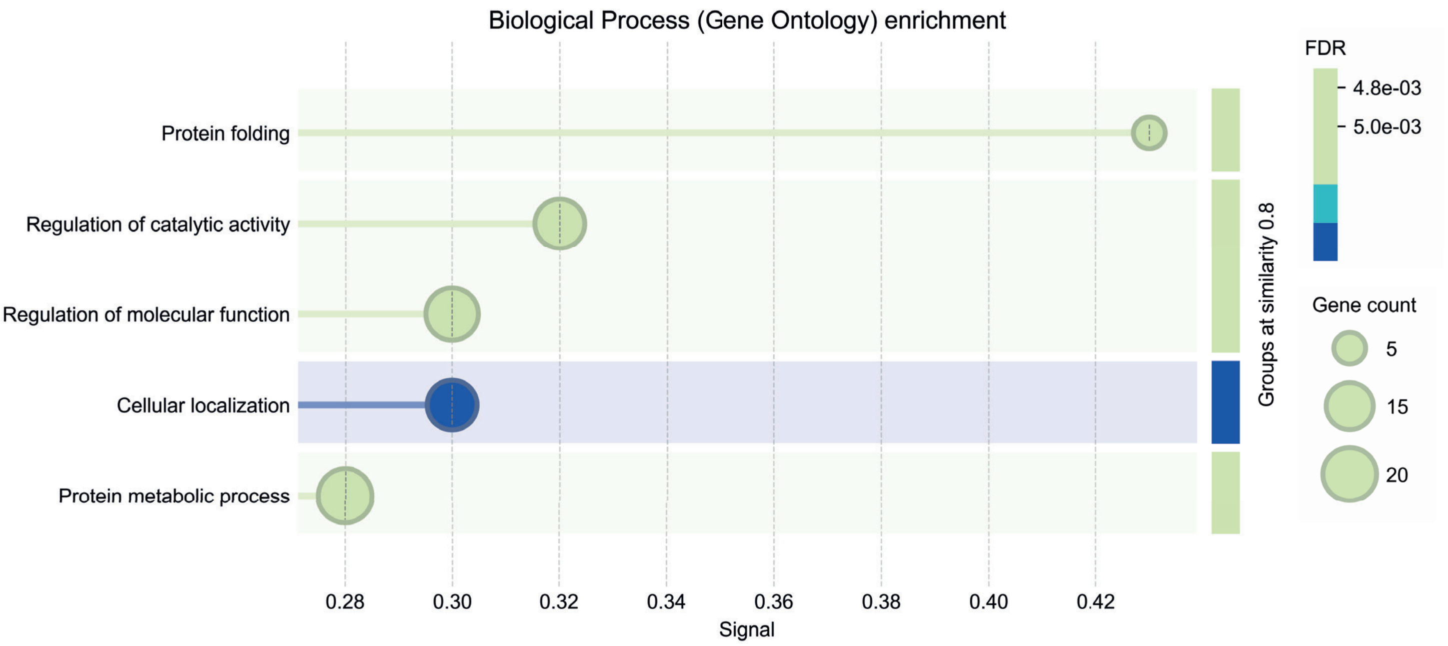


Suppl. Fig. 8

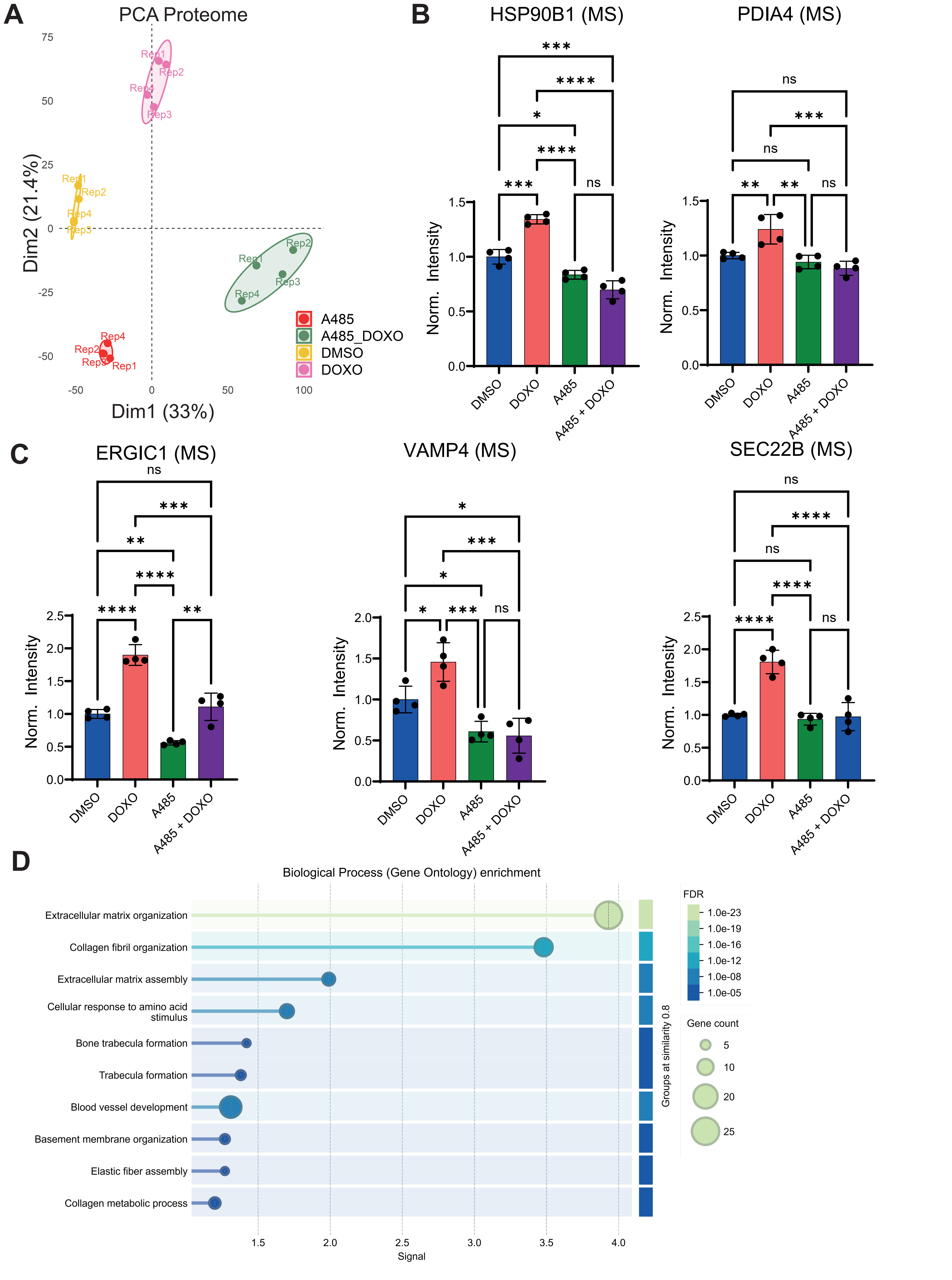
A
PCA Proteome
Dim1 (33%)
Dim2 (21.4%)
A485
A485_DOXO
DMSO
DOXO
B
HSP90B1 (MS)
PDIA4 (MS)
Norm. Intensity
DMSO
DOXO
A485
A485 + DOXO
C
ERGIC1 (MS)
VAMP4 (MS)
SEC22B (MS)
D
Biological Process (Gene Ontology) enrichment
Extracellular matrix organization
Collagen fibril organization
Extracellular matrix assembly
Cellular response to amino acid stimulus
Bone trabecula formation
Trabecula formation
Blood vessel development
Basement membrane organization
Elastic fiber assembly
Collagen metabolic process
Signal
Groups at similarity 0.8
FDR
1.0e-23
1.0e-19
1.0e-16
1.0e-12
1.0e-08
1.0e-05
Gene count
5
10
20
25

Suppl. Fig. 9
